# QKI ensures splicing fidelity during cardiogenesis by engaging the U6 tri-snRNP to activate splicing at weak 5ʹ splice sites

**DOI:** 10.1101/2025.09.04.674271

**Authors:** Maureen V Akinyi, Wenjie Yao, Jakub Zeman, Clara Hipp, Deniz Bartsch, Laurence Heaven, Charlotte A Le Roux, Anne C Starner, Fei Yuan, Mason D Bartels, Fangyu Zhao, Hang Le Ha, Riddhi Sharma, Bhumika Choudhary, Josep Biayna, Ralf P Brandes, Gabrijela Dumbović, Arjun Ray, Ilka Wittig, Christian Muench, Michael Sattler, Eric L Van Nostrand, Leo Kurian

## Abstract

During organogenesis, precise pre-mRNA splicing is essential to assemble tissue architecture. Many developmentally essential exons bear weak 5′ splice sites (5′SS) yet are spliced with high precision, implying unknown yet active splicing fidelity mechanisms. By combining transcriptome and alternative splicing profiling with temporal eCLIP mapping of RNA interactions across development, we identify the RNA-binding protein QKI as an essential direct regulator of splicing fidelity in key cardiac transcripts. Although QKI is dispensable for cardiac specification, its loss disrupts sarcomere assembly despite intact expression of sarcomere mRNAs through exon skipping and nuclear retention of mis-spliced RNAs. QKI-dependent exons in essential cardiac genes have weak 5′SS and frequently show poor complementarity with U6 snRNA. We show that QKI directly interacts with U6 snRNA using an overlapping interface to its traditional intronic binding activity, securing U4/U6·U5 tri-snRNP to ensure splicing fidelity. Thus, QKI exemplifies how context-aware RBPs enforce splicing fidelity at structurally vulnerable splice sites during organogenesis.

## MAIN

Pre-mRNA splicing is a fundamental step in gene expression, enabling the programmed and precise reconstitution of the segmented coding information in eukaryotic DNA^1–4^. During embryonic development, global rewiring of alternative splicing networks is critical for precise cell fate decisions across organs such as the brain, heart, liver, and germline, and disruption of these programs underlies a broad spectrum of congenital and malignant diseases^5–7^. Conserved alternative splicing programs are especially prominent in the brain and heart, underscoring a central role for cell-type-specific, regulated splicing in organogenesis^5–17^. Alternative splicing is pervasive, occurring in over 90% of multi-exonic human genes, and generates extensive isoform diversity^1, 18^. This process not only reshapes coding sequences to generate protein isoforms but also remodels untranslated regions to influence transcript localization, translation, and stability^2^. Splicing is governed by highly complex and context-specific regulatory logic, and while its core mechanisms are well established, a major challenge in the field remains in defining the rules that shape cell-type-specific splicing architecture and predict splicing outcomes in physiologically relevant contexts^19–21^.

Successful splicing of an intron depends on the accurate recognition of 5ʹ and 3ʹ splice sites by the spliceosome^19, 22–24^. Pre-mRNA splicing is orchestrated by the spliceosome, a dynamic ribonucleoprotein machinery comprising over 300 proteins and five uridine-rich snRNAs (U1, U2, U4, U5, U6), which sequentially assemble on pre-mRNA to catalyze intron removal^24^. Splicing outcomes are determined on a case-by-case basis by the interplay of cis-acting RNA sequence elements and structures within exons and introns, and trans-acting factors—including snRNAs and RNA-binding proteins (RBPs), as well as chromatin architecture and transcriptional dynamics^4,20, 25–27^.

While splice site recognition is remarkably robust, allowing precise exon selection across vast transcriptomic complexity, it is particularly reliant on accurate recognition of the 5ʹ splice site (5ʹSS)̶the entry point for spliceosome assembly. A pivotal step in this process is the engagement of the pre-assembled U4/U6.U5 tri-snRNP with splice sites to form the catalytically competent spliceosome^22, 28^. Although U1 initiates 5ʹSS recognition, it is U6 snRNA that ultimately pairs with the 5ʹSS to trigger catalysis^22^. Strikingly, a substantial fraction of human exons harbor weak, non-consensus 5ʹSS with limited complementarity to spliceosomal snRNAs^29, 30^. Weak splice site scores have long been associated with alternative splicing^31^, yet high precision of alternative splicing plays a disproportionally critical role in brain and heart development^6^. This paradox points to the existence of specialized, context-aware mechanisms that ensure splicing fidelity when core sequence signals are insufficient, though the molecular identity and logic of such mechanisms have remained elusive^21, 22, 30, 32^. Deciphering how such structurally vulnerable splice sites are reliably recognized is central to understanding the regulatory architecture of gene expression during embryonic development^5, 7^.

One RBP that has been linked to control of exon inclusion, during heart and brain development, is QKI^6^. Recently, an elegant cross-species developmental transcriptome analysis revealed correlative enrichment of QKI-binding motifs, particularly the *ACTAAC* variant, near cassette exons with weak 5ʹSS^6^. These QKI binding motifs exhibit positional distribution in tissue-specific transcripts with notable enrichment in cardiac transcripts^6, 33^. However, the mechanistic relevance of this pattern remains unknown. In line with these observations, QKI loss in both human and mouse models leads to severe defects in heart development and function as well as alteration of the alternative splicing landscape^34, 35^. QKI is broadly implicated in cell-state transitions and pathology - regulating neuronal myelination, hematopoietic differentiation, and cardiac sarcomere formation, and, in cancer, rewiring stress-granule–linked mRNA metabolism^36–41^. Intriguingly, despite being ubiquitously expressed, QKI appears to regulate splicing in a highly context-dependent manner. However, how QKI maintains the integrity of cardiac splicing architecture has yet to be determined, and how such a broadly expressed RBP recognizing a widely distributed motif exerts selective control over organ-specific splicing programs during embryogenesis remains unresolved.

Here, we uncover a mechanism by which QKI ensures exon inclusion in essential cardiac genes during cardiac commitment of human pluripotent stem cells. We reveal that QKI-deficient cardiomyocytes express the full complement of lineage-specific mRNAs as wild-type counterparts yet fail to assemble the contractility apparatus due to mis-splicing and nuclear retention of key sarcomeric transcripts. Using stage-resolved transcriptomics and eCLIP-seq, we show that QKI dynamically reshapes its RNA interactome, progressively interacting with transcripts encoding the contractility apparatus and playing a critical role in maintaining splicing architecture in terminally differentiated cardiomyocytes. Mechanistically, we show that QKI not only binds intronic motifs near weak 5′SS but also directly engages the U6 snRNA, likely providing necessary reinforcement of tri-snRNP engagement at weak splice sites where splicing fidelity is inherently compromised. Biochemical and functional analyses reveal that QKI engages both pre-mRNA near weak 5ʹSS and U6 via overlapping interfaces and interacts with core tri-snRNP components, integrating splice site recognition with spliceosome assembly to enforce splicing fidelity in transcripts essential for organ function during development. These findings illuminate a previously unknown mechanism of how splicing accuracy is maintained during organogenesis and offer a broader framework for how ubiquitously expressed RBPs, despite recognizing widespread motifs, selectively program splicing to preserve gene expression fidelity during embryonic development.

## RESULTS

### QKI uncouples cardiac fate specification from sarcomere formation and contractile function

Although QKI has been implicated in both early lineage specification^35^, as well as neuronal^41, 42^ and cardiac development^34, 36^, its function appears to be highly context dependent. This is especially notable in the developing heart, given that QKI is broadly expressed as confirmed at both RNA and protein levels (**Extended Data Fig. 1a-c**) and nuclear localized (**Extended Data Fig. 1d,e)** throughout cardiac specification of human embryonic stem cells (hESCs).

To systematically evaluate the developmental dependencies and define the cardiac cell type specification window in which QKI becomes essential, we generated hESC clones lacking all QKI isoforms. CRISPR-Cas9-mediated targeting of the constitutive exon 3 (**Extended Data Fig. 2a)** resulted in the complete loss of all QKI protein isoforms, yielding three independent homozygous QKI knockout (QKI KO) hESC clones (**Extended Data Fig. 2b,c**). Despite its high expression levels in pluripotency, loss of QKI did not affect self-renewal of hESCs or the expression of key pluripotency factors, except for a reduction in SOX2 levels (**Extended Data Fig. 2d-h)**. Unlike previously reported^34, 35^, in our hands, QKI loss did not affect germ layer competence of hESCs (**Extended Data Fig. 3)**. QKI-KOs efficiently and robustly differentiated into three primary germ layers: mesoderm (**Extended Data Fig. 3a)**, endoderm (**Extended Data Fig. 3b)**, and ectoderm (**Extended Data Fig. 3c)**, evaluated using the germ layer commitment protocol we previously reported^43, 44^ and profiling the indicated lineage-specific markers in all three QKI KO clones.

Next, we assessed the cardiac differentiation potential of QKI KO hESCs using a robust directed differentiation protocol we previously established^43–45^ (**Fig. 1a**). This approach applies a WNT-BMP concentration gradient (“cardiac corridor”) to buffer cell line-specific morphogen requirements and systematically probe the capacity of a given perturbation to commit to the cardiac lineage, followed by WNT inhibition to drive efficient cardiac specification. Consistent with previous findings^34^, we observed that QKI KO cardiomyocytes lacked sarcomere architecture, hallmarks of cardiac function (**Fig. 1b-g**). Previous studies linked this loss of proper cardiomyocyte specification to a role for QKI in mesoderm commitment and in early cardiac commitment, with one reporting impaired germ layer formation^35^ and another suggesting defective cardiac induction^34^. However, we observed in our model that QKI loss did not impair cardiac specification, as all three KO clones showed robust expression of core cardiac transcription factors such as NKX2.5 (**Fig. 1c**), GATA6 (**Fig. 1d),** and MEF2C (**Fig. 1e**). Instead, we observed that QKI KO caused a shift in morphogen requirement to commit to cardiac mesoderm, as KOs differentiated optimally at lower WNT levels (**Fig. 1a**), potentially due to reduced SOX2 expression in pluripotency, since downregulation of SOX2 is a foundational step in mesoderm commitment driven by coordinated WNT–BMP signaling^45^. As QKI-deficient hESCs retain pluripotency (**Extended Data Fig. 2**), differentiate into all three germ layers (**Extended Data Fig. 3**), and efficiently generate cardiomyocytes when morphogen levels are adjusted, we hypothesize that the previously reported mesodermal defects may reflect indirect effects due to altered morphogen sensitivity upon QKI loss.

**FIGURE 1.**
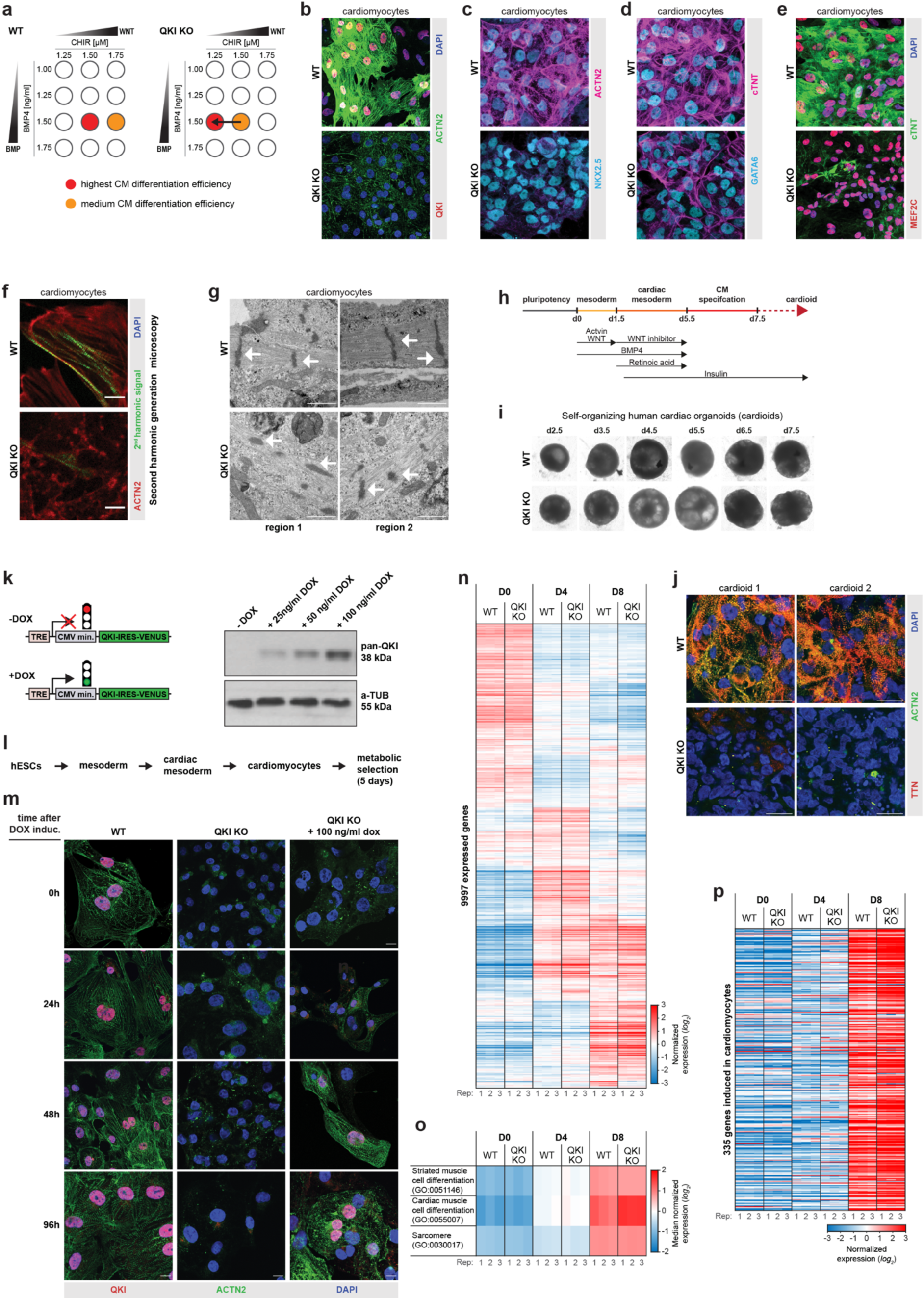
| QKI uncouples cardiac fate specification from sarcomere formation and contractile function. **a,** Schematic summarizing the cardiac differentiation efficiency along the WNT-BMP morphogen gradient (“cardiac corridor”) for wild-type (WT) and QKI knockout (KO) hESCs. QKI KO cells efficiently generated cardiomyocytes, albeit with shifted optimal WNT requirements. (n = 3 independent biological replicates). **b-e**, QKI KO cardiomyocytes show robust expression of core cardiac transcription factors but severely disrupted sarcomere organization. Immunofluorescence staining for cardiomyocyte markers (**b**) ACTN2 and cardiac troponin T (cTnT) and indicated cardiac transcription factors (**c**) NKX2.5, (**d**) GATA6, and (**e**) MEF2C in WT and QKI KO cardiomyocytes. Scale bars, 20 μm. **f**, Second harmonic generation (SHG) imaging of sarcomere structure in WT and QKI KO cardiomyocytes reveals lack of organized contractile structures in QKI KO. Scale bar, 10 µm. **g**, Transmission electron microscopy (TEM) showing sarcomeric architecture in WT and QKI KO cardiomyocytes at day 8. QKI KO cardiomyocytes lack defined Z-lines and regular sarcomere units. Scale bar, 500 nm. **h**, Schematic of cardioid generation method. **i**, Representative brightfield images of WT and QKI KO cardioids on day 8 showing comparable size and morphology. **j**, Immunofluorescence staining for TTN and ACTN2 in WT and QKI KO cardioids at day 10 shows disorganized sarcomere structure in QKI KO organoids. Scale bar, 20 µm. **k**, Illustration of the strategy used to re-express QKI in QKI-KO cardiomyocytes using PiggyBac-based strategy driving QKI-mVENUS using doxycycline (DOX) inducible promoter. Right: immunoblot confirming QKI re-expression at WT levels. **l**. Schematic of the generation and metabolic maturation of cardiomyocytes to deplete any carry over of proliferating cells post differentiation. **m**. Rescue of sarcomere assembly following DOX-induced QKI re-expression in QKI KO cardiomyocytes. Immunostaining shows progressive restoration of sarcomere structure upon QKI re-expression in QKI KO cardiomyocytes. Cells were differentiated without DOX, metabolically selected, replated and induced with DOX for 96 hours. Sarcomeres reappear 24-48 hours post-induction. Scale bar, 20 µm. **n**, Heatmap showing transcriptome-wide expression profiles in wild-type and QKI KO cells (average FPM across 3 replicates) at key stages of directed cardiac differentiation—pluripotency (D0), cardiac progenitor (D4), and cardiomyocyte (D8). Despite loss of QKI, global transcriptional programs proceed largely unchanged between WT and QKI KOs in the indicated stages. **o**, QKI loss does not impair functional gene activation during cardiac differentiation. Gene Ontology (GO) enrichment analysis of differentially expressed genes in wild-type and QKI KO cells at key stages of differentiation reveals near-identical, stage-specific activation of gene programs associated with cardiomyocyte specification, sarcomere assembly, and cardiac contractility. **p**. Heatmap showing that genes specifically induced during cardiomyocyte differentiation exhibit near-identical expression dynamics in wild-type and QKI knockout cells at day 8, indicating intact transcriptional activation of cardiac identity programs despite QKI loss. (n = 3 in all experiments unless otherwise stated).

Intriguingly, QKI KO cardiomyocytes showed severe depletion of key sarcomeric proteins, including ACTN2 and cTNT (**Fig. 1b–e**). To rule out potential loss of antibody epitope due to potential splicing defects in QKI-KOs, we confirmed the absence of organized sarcomeres using two orthogonal, label-free approaches: second harmonic generation (SHG) microscopy (**Fig. 1f**) and transmission electron microscopy (TEM) (**Fig. 1g**). Both approaches revealed fragmented, sarcomere-like structures with severely compromised architecture, confirming a failure to assemble functional sarcomeres in the absence of QKI. These findings were further validated using a cardiac organoid (cardioid) model^46^ (**Fig. 1h**). QKI-KOs efficiently generated cardioids comparable in size to their isogenic wild-type counterparts (**Fig. 1i**). Consistent with findings from the 2D differentiation model, QKI-KO cardioids exhibited a lack of organized and functional sarcomere architecture as measured by expression patterns of TTN and ACTN2 in Day 8 cardioids evaluated by immunofluorescence (**Fig. 1j**). Together, these data reveal that while QKI is dispensable for cardiomyocyte specification, it is essential for sarcomere assembly and architecture.

We reasoned that if QKI knockout cells are indeed fully specified cardiomyocytes lacking sarcomeres, then reintroducing QKI after cardiac differentiation should rapidly restore sarcomere assembly. This approach would provide definitive evidence that differentiated QKI KOs are indeed cardiomyocytes and directly demonstrate that QKI decouples cardiac specification from functional maturation. To test this, we generated a PiggyBac-based system to introduce a doxycycline (DOX)-inducible *QKI–mVENUS* transgene into QKI-KO hESCs (**Fig. 1k**). We used the QKI-5 isoform for this purpose, as it is the predominant QKI isoform expressed during cardiac development and is exclusively localized to the nucleus. These engineered cells were differentiated into cardiomyocytes without QKI induction and then replated in glucose-free medium supplemented with fatty acids and thyroid hormone to promote metabolic maturation (**Fig. 1l**). This 10-day protocol selectively enriches cardiomyocytes by eliminating proliferative cells. We then initiated a 96-hour QKI re-expression time course via DOX treatment. Sarcomeres were absent in QKI-KO cardiomyocytes but began to reappear within 24–48 hours of Dox-induction, which restores QKI to wildtype levels (**Fig. 1m**). These results confirm that QKI is dispensable for cardiomyocyte specification but essential for sarcomere assembly, functionally uncoupling cardiac identity from contractility.

Finally, we hypothesized that if QKI loss does not impair cardiac specification, QKI KO and wild-type (WT) cells should display similar gene expression profiles throughout early differentiation, with differences, if any, emerging only at the cardiomyocyte stage. To test this, we performed whole-transcriptome sequencing at defined stages of cardiac differentiation: day 0 (pluripotency), day 4 (cardiac progenitors), and day 8 (cardiomyocytes). Analysis of 9,997 reliably expressed genes revealed highly similar transcriptional dynamics between QKI KO and WT cells at all stages (**Fig. 1n, Supplementary Table 1**). Specifically, gene expression dynamics of hallmark cardiomyocyte programs and genes essential for sarcomere formation showed nearly identical activation between QKI KO and WT cells, illustrated by gene ontology (GO) based functional enrichment analysis (**Fig. 1o**). Remarkably, QKI KO cardiomyocytes even showed robust expression of genes involved in muscle differentiation and sarcomere assembly, including considering the 335 genes linked to sarcomere organization and contractility, at levels comparable to their WT counterparts (**Fig. 1p**). These findings reveal a striking disconnect between transcriptional identity and structural outcome, as QKI KO cardiomyocytes express the full complement of contractile genes but fail to assemble functional sarcomeres (**Fig. 1b-f**). Collectively, these findings demonstrate that QKI uncouples cardiomyocyte identity from contractility, serving as a critical regulator that separates cell fate specification from functional acquisition during heart development.

### Stage-resolved eCLIP maps transcriptome-wide QKI binding landscapes during human cardiac specification

Our observation that QKI loss selectively disrupts sarcomere formation and contractility without affecting cardiac specification reveals a disconnect between its expression pattern and stage-specific function. While prior efforts have explored QKI-dependent events based on the presence of QKI motifs^47, 48^, the fact that QKI is expressed throughout cardiac differentiation (**Extended Data Fig. 1a-c**) suggests that the definition of its mechanistic role in heart development requires experimental stage-resolved mapping of QKI–RNA interactions. We therefore performed enhanced crosslinking immunoprecipitation followed by sequencing (eCLIP-seq)^49, 50^ across key stages of human cardiac commitment, namely pluripotency, cardiac progenitors, and cardiomyocytes, to chart the time-resolved RNA interaction landscape of QKI (**Fig. 2a, Extended Data Fig. 4a**).

**FIGURE 2.**
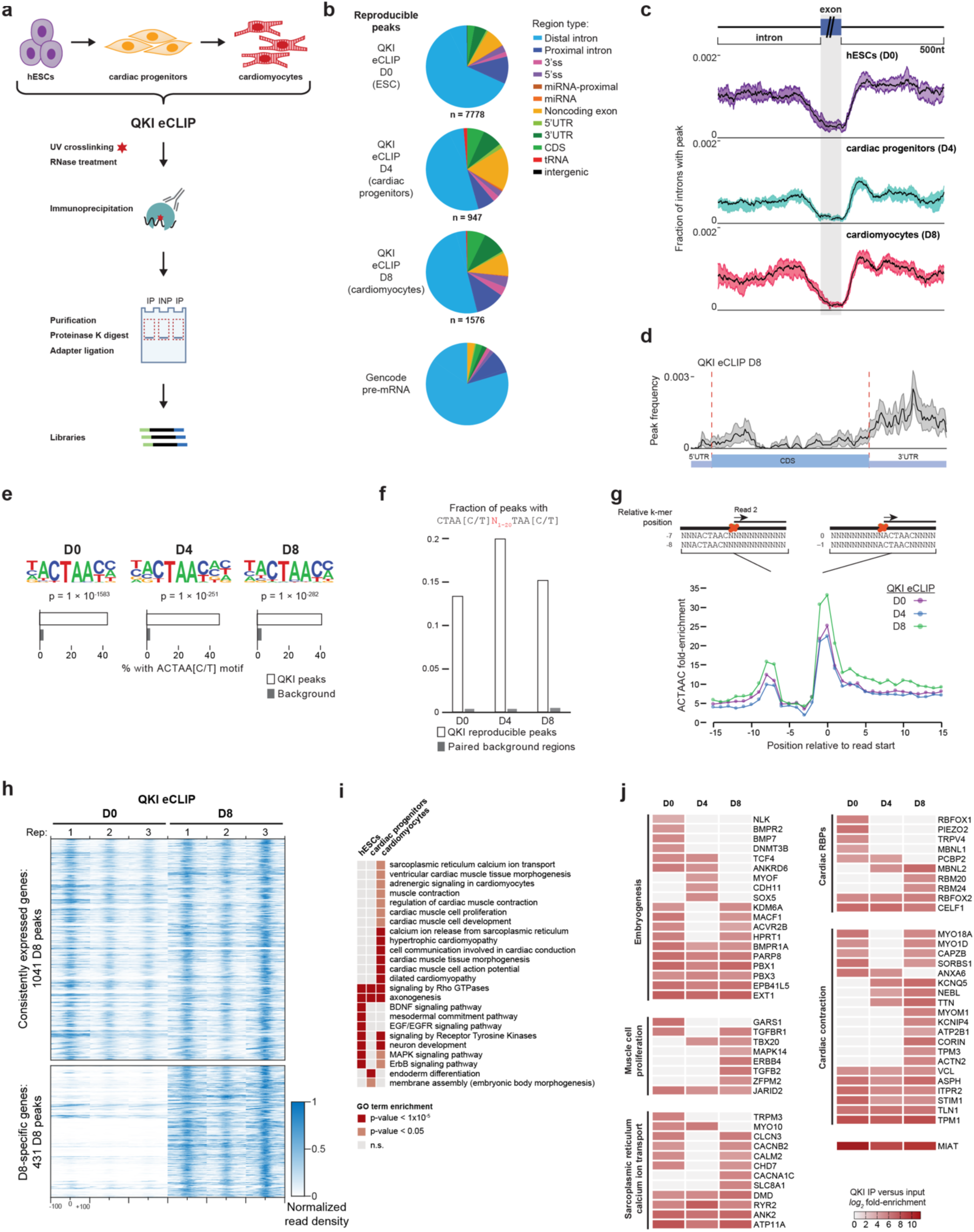
| Stage-resolved eCLIP-seq maps transcriptome-wide QKI binding landscapes during human cardiac specification. **a**, Schematic of stage-resolved eCLIP-seq across human cardiac differentiation from pluripotency (D0), cardiac progenitors (D4), to cardiomyocytes (D8). **b**, Total number of reproducible, significantly enriched QKI binding eCLIP peaks per stage normalized against the paired SMInput (n=3 biological replicates at each stage of differentiation, fold change ≥ 2; *q*-value ≤ 0.05). Pie chart shows the genomic distribution of QKI peaks across introns, 3ʹUTRs, coding regions, and other regions across all stages versus total nucleotides covered in Gencode (v19) transcripts. **c**, Metagene plot shows intronic QKI eCLIP peak distribution for significant peaks (*p* ≤ 10^-3^, fold-enrichment ≥ 8) from example individual replicate eCLIP, showing enrichment near 5ʹSS in all indicated time points of cardiac differentiation. **d,** Metagene plot for cardiomyocyte QKI eCLIP peaks along a meta-mRNA shows enrichment in the 3’UTR. **e**, Top enriched QKI-binding motif (ACTAAY) and bipartite consensus (ACTAA[C/T]N₁–₂₀TAA[C/T]) derived from HOMER analysis of IDR eCLIP peaks, with fold-enrichment above background for the indicated time points **f**, Percent of peaks at each stage containing the canonical bipartite QKI motif. **g**, Reverse transcription stop-site (RT-stop) analysis across QKI-bound motifs reveals increased termination at -1, 0 and -7/-8 positions relative to QKI motif locations. **h**, Peak read density in QKI eCLIP (normalized across both timepoints) for QKI eCLIP peaks identified in D8, separated by their location in (bottom) D8-specific genes (D8 expression more than 5-fold increased from D0) or (top) other genes. **i**, GO enrichment analysis of QKI-bound mRNAs at each stage, showing a shift from early neuronal and signal transduction components (D0, D4) to sarcomere and contraction-related categories at D8. **j**, Heatmap of representative QKI-bound transcripts grouped by function. At D8, QKI selectively binds mRNAs encoding cardiac-relevant RBPs such as *RBM20, RBM24, RBFOX2, MBNL1/2,* as well as sarcomeric and calcium-handling proteins, including *TTN, RYR2, ACTN2, TPM1, NEBL*, and *CAMK2D*.

After removal of PCR duplicates and normalization to size-matched input controls, we generated high-confidence, nucleotide-resolution QKI binding maps from three independent biological replicates (**Extended Data Fig. 4b**). Using a custom adaptation of the Irreproducible Discovery Rate framework^51^ to incorporate three replicates, we identified 7,778 reproducible QKI peaks at D0 (pluripotency), 947 at D4 (cardiac progenitors), and 1,576 at D8 (cardiomyocytes) (**Fig. 2b**).

Across all timepoints, ∼74% of QKI binding events localized to introns (**Fig. 2b, Extended Data Fig. 4b**), with a particular enrichment in 5ʹ and 3ʹ splice sites as well as adjacent proximal intronic regions, consistent with roles in splicing regulation (**Fig. 2c**). A smaller fraction of peaks was observed in 3’ UTRs (**Fig. 2b,d, Extended Data Fig. 4b**), in line with additional functions reported for QKI in mRNA stability (particularly in neuronal lineages). Motif enrichment analysis using HOMER revealed a dominant ACTAAY motif enriched >15-fold above background and present in >40% of peaks (**Fig. 2e-g**), matching the known in vitro QKI recognition sequence. Furthermore, a bipartite consensus motif (ACTAA[C/T]N₁–₂₀TAA[C/T]) previously shown^47, 48^ to mediate high-affinity QKI binding was found in 13-20% of peaks (>33-fold enrichment) (**Fig. 2f**), reinforcing the specificity and quality of our datasets. Reverse transcription stop-site analysis relative to QKI motifs revealed stereotypical pileups at position 0 and -1, with secondary enrichment at -7/-8 (**Fig. 2g**), consistent with reverse transcription termination at crosslinked direct QKI–RNA contacts.

QKI is expressed throughout cardiac differentiation (**Extended Data Fig. 1a-c**), and clustering analysis indicated a conserved QKI interactome network, with all QKI samples—including in-house replicates as well as prior experiments from the ENCODE consortium^50, 52^—forming a distinct cluster separate from unrelated eCLIPs such as IGF2BP1–3 and RBFOX2 in hESCs (**Extended Data Fig. 4c**). However, considering the cardiomyocyte interactome, we observed two distinct sets of targets: 72% of peaks were found in genes consistently expressed between stem cells and cardiomyocytes, and those were consistently QKI-bound as early as stem cells (**Fig. 2h, Extended Data Fig. 4d**). In contrast, the 28% of peaks found in cardiomyocyte-specific genes (>5-fold induced in cardiomyocytes) were largely only QKI-bound in cardiomyocytes (**Fig. 2h, Extended Data Fig. 4d**). Thus, underlying gene expression change enables a dramatic retuning of the QKI regulatory landscape as cardiogenesis proceeds, maintaining a broad network present in pluripotent cells while newly including a more focused cardiac-specific program post-specification.

Although QKI binds broadly in pluripotent hESCs (D0) and cardiac progenitors (D4), these targets comprised a heterogeneous set of developmental regulators without a unifying lineage identity (**Fig. 2b,i,j**). Notably, functional annotation using gene ontology (GO term) analysis at early stages revealed an unexpected enrichment for neuronal genes, but not cardiac-associated transcripts (**Fig. 2i,j**). This early binding profile did not translate into phenotypic defects (**Fig. 1a, Extended Data Fig. 2, 3**), aligning with the lack of observable requirement for QKI during early cardiac induction. In stark contrast, QKI binding in cardiomyocytes (D8) showed a pronounced and functionally coherent enrichment for mRNAs encoding key components of the contractile apparatus and calcium signaling pathways (**Fig. 2i,j**). These included core sarcomeric genes such as *TTN, NEBL, ACTN2, TPM3, TLN1*, and *STIM1*, cell cycle regulators such as *NIN*, as well as central regulators of calcium handling and excitation–contraction coupling such as *RYR2, ATP11A, SLC8A1, CAMK2D*, and *CACNA1C* (**Fig. 2j**). GO enrichment analysis of D8-bound transcripts underscored this transition, with top categories including sarcomere organization, muscle contraction, actin filament-based processes, and regulation of cytosolic calcium ion concentration (**Fig. 2i**).

Together, these data provide the first stage-resolved, transcriptome-wide binding maps of QKI during human cardiac specification, offering a comprehensive view of its temporal target landscape. Surprisingly, they reveal that the QKI RNA interactome is dynamically rewired during cardiac differentiation, with a pronounced shift toward transcripts that govern the contractile machinery in terminally differentiated cardiomyocytes.

### QKI ensures splicing fidelity in essential cardiac genes

Our observation that QKI loss has a significant impact on cardiomyocyte function but not gene-level RNA expression (**Fig. 1n-p**) is consistent with prior findings that QKI has a particular impact on alternative splicing in cardiomyocytes^34, 36^. To identify splicing events regulated by QKI during cardiac differentiation, paired with our direct targets identified by eCLIP, we performed differential splicing analysis using rMATS^53^ from our RNA-seq data profiling wild-type and QKI KO cells at matched timepoints (day 0, 4, and 8) (**Fig. 3a-c**). Events were required to have at least 10 junction-spanning reads and meet a false discovery rate threshold of ≤ 0.01, ensuring high-confidence detection of QKI-dependent exon inclusion and exclusion. The most prominent splicing changes were in the inclusion and exclusion of cassette exons, although other alternative splicing events were also present to a lesser extent (**Fig. 3a**). In total, we observed hundreds of significantly QKI knockout-included (255 (D0), 132 (D4), and 469 (D8)) or knockout-excluded (362 (D0), 129 (D4), and 351 (D8)) exons at each timepoint (**Fig. 3b,c, Supplementary Table 2**).

**FIGURE 3.**
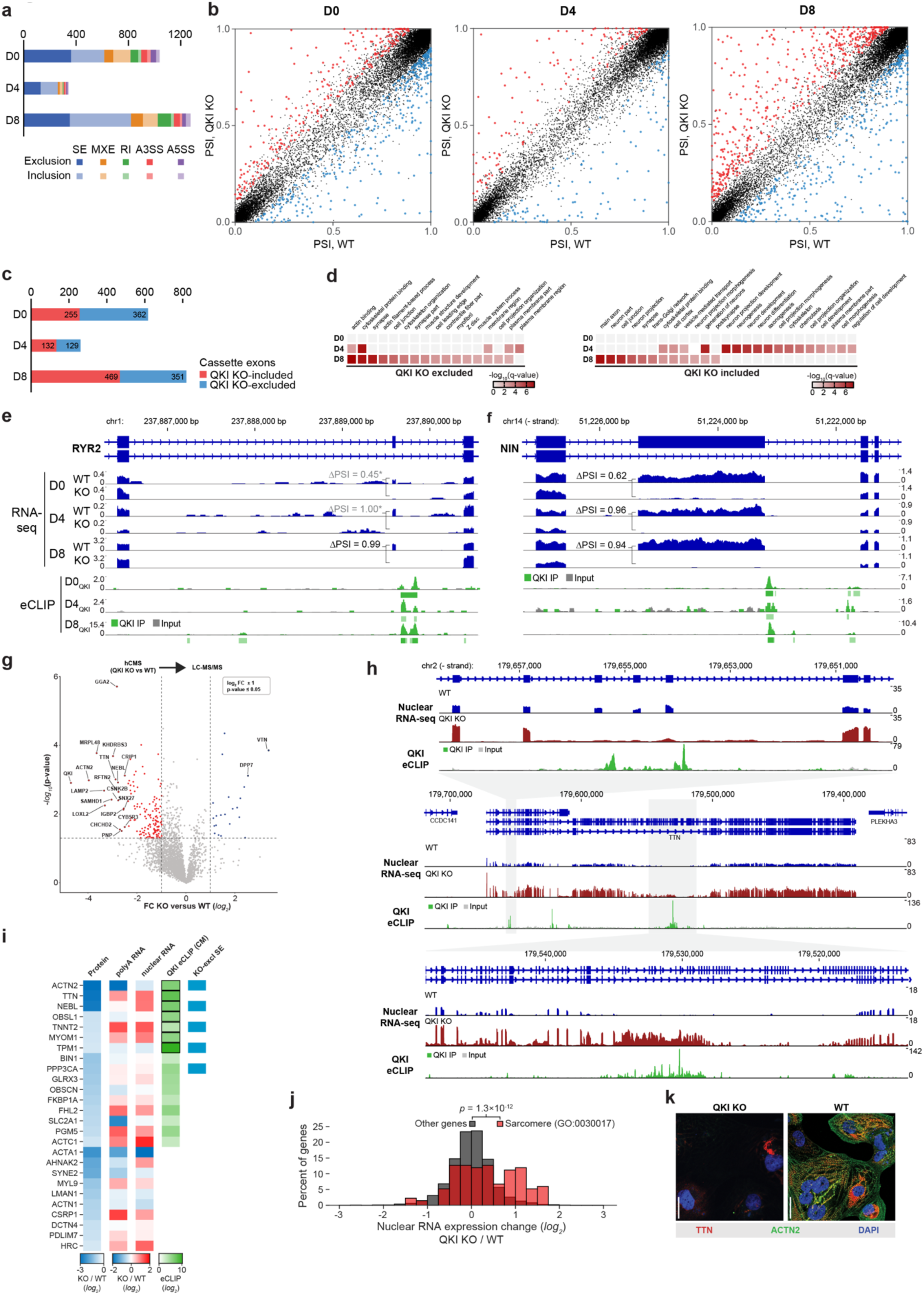
| QKI loss leads to altered RNA splicing, RNA nuclear retention, and protein abundance of sarcomere genes. **a**, Bar plot showing the number of significant splicing events (FDR ≤ 0.01, ΔPSI ≥ 10%) identified in QKI KOs across D0, D4, and D8, categorized into skipped exon (SE), mutually exclusive exon (MXE), retained intron (RI), alternative 3ʹ (A3SS), and 5ʹ splice sites (A5SS). **b**, Dot plots highlighting differentially included (red, ΔΨ > 0) and excluded (blue, ΔΨ < 0) cassette exons in QKI KO versus WT across differentiation stages. **c**, Bar plots showing the number of significant alternative splicing events detected per timepoint in QKI knockout. **d**, Gene Ontology (GO) term enrichment analysis of cassette exon (left) exclusion and (right) inclusion events in QKI KO events across D0, D4, and D8 stages indicates D8 QKI-dependent exon inclusion at D8 enriched in contractile and sarcomeric programs while exclusion events are enriched in neuronal and migration-related genes. **e**,**f**, Read density of QKI eCLIP and RNA-seq at alternative exons in (**e**) *RYR2* and (**f**) *NIN* with significant exclusion in QKI KO versus WT indicates QKI binding proximal to the alternative exon. Grey PSI (Percent Spliced In) indicates quantitation from less than 10 junction-spanning reads. **g**, Volcano plot of quantitative whole cell LC-MS/MS proteomics (WT vs. QKI KO cardiomyocytes, *n = 3* biologically independent replicates) indicates depletion of key sarcomeric proteins among QKI targets. *P* values are calculated using Student’s *t* test; biological replicates *n* = 3 **h**, Read density of QKI eCLIP and nuclear RNA-seq at *TTN* in WT and QKI KO cardiomyocytes indicates QKI binding near (top) 5ʹSS of exons with QKI-dependent inclusion and (bottom) nuclear accumulation of mis-spliced *TTN* transcripts in QKI KO cardiomyocytes. **i,** Shown are all sarcomere genes with differential (*log_2_*(FC) ≤ -0.5) protein abundance in QKI KO versus WT, along with fold-change observed in proteomic, whole-cell polyA RNA-seq, and nuclear total RNA-seq, along with indication of the presence of eCLIP peaks and QKI KO-dependent splicing changes. eCLIP indicates the maximum fold-enrichment across 3 biological replicates, with reproducible peaks across all replicates indicated with black outline. **j**, Histogram indicates the increase in nuclear versus total RNA abundance in QKI KO versus WT for sarcomere genes. Significance was determined by Kolmogorov-Smirnov test. **k**, Representative immunofluorescence images showing near-complete loss of TTN and ACTN2 protein in QKI KO cardiomyocytes, despite transcriptional induction. Scale bar, 10 µm.

Although QKI binds broadly in pluripotent hESCs (D0) with significant splicing changes upon its loss, affected transcripts generally lacked coherent functional enrichment (**Fig. 3d**). In cardiac progenitors (D4), QKI-KO-induced exon inclusions were unexpectedly enriched in neuronal genes, suggesting a role for QKI in repressing non-cardiac exons (**Fig. 3d**). In contrast, QKI-dependent exon inclusion (i.e., KO-excluded exons) at the cardiomyocyte stage (D8) predominantly affected transcripts essential for cardiac structure and contractility (**Fig. 3d**), highlighting a central role of QKI in ensuring proper exon inclusion in genes encoding the cardiac contractility apparatus.

Our QKI eCLIP data enabled us to identify directly regulated QKI-dependent splicing events based on concurrent QKI binding. Several events observed in our dataset provide empirical transcriptome-wide validation of direct QKI binding at QKI-dependent splicing events previously observed in mouse cardiomyocytes or whole heart tissue, such as *ACTN2, BIN1, RYR2, NEBL, AKAP9*, *RBFOX2,* and *TTN*^34, 36^ or in neuronal cells during development, including *NIN* (a key regulator of the cell cycle)^54, 55^ (**Fig. 3e,f, Extended Data Fig. 5a-c**). Extending these observations, our data suggest that QKI directly controls exon inclusion in a broad repertoire of cardiac-critical transcripts, including actin capping factor *CAPZB, TPM1, OBSL1, and TNNT2* (**Extended Data Fig. 5d**). To test whether this regulatory logic is generalizable beyond cardiomyocytes, we constructed minigene reporters spanning five representative QKI-bound, differentially included exons and transfected them into HEK293T cells. Four out of five constructs faithfully recapitulated the endogenous QKI-dependent splicing response (**Extended Data Fig. 5e**), demonstrating that QKI binding alone is sufficient to drive exon inclusion of exons in genes critical for cardiomyocyte function, even outside the cardiac context. These results underscore that the role of QKI in ensuring exon inclusion at cardiac-relevant exons reflects a more universal regulatory principle.

### QKI loss leads nuclear sequestration of mis-spliced sarcomeric transcripts, resulting in depletion of essential sarcomeric proteins

Although QKI KO cardiomyocytes express cardiac transcription factors and sarcomeric genes at levels comparable to wild type, they fail to assemble functional sarcomeres (**Fig. 1**), suggesting a disconnect between transcript abundance and protein output. To explore how QKI binding and alternative splicing control could explain this disconnect, we performed unbiased whole-cell proteomic profiling using LC-MS/MS. As expected, QKI was the most depleted protein in knockout cells (**Fig. 3e**). Aligning with the inability to assemble sarcomeres, we observed that among the top 10 most depleted proteins were three core proteins essential for cardiomyocyte structure and contractile function: TTN (Titin), NEBL (Nebulin), and ACTN2 (Alpha-actinin-2) (**Fig. 3g-i**). Inspection of all three revealed not only significant QKI binding in cardiomyocyte eCLIP, but also a variety of mis-splicing patterns in QKI knockout cells (**Fig. 3h, Extended Data Fig. 6a,b**). Although *ACTN2* RNA was decreased (4.5-fold) in QKI knockout, *NEBL* RNA was 1.7-fold increased (**Fig. 3i, Extended Data Fig. 6a,b**), indicating a dichotomy between RNA and protein expression. Extending this to all sarcomere proteins (GO:0030017) yielded similar results, with a general lack of correlation between RNA and protein changes (**Extended Data Fig. 6c**).

This disconnect prompted us to investigate whether QKI-dependent splicing defects might impair the maturation and export of mRNA to the cytoplasm. To systematically test this, we performed matched nuclear and whole-cell total RNA-seq in WT and QKI KO cardiomyocytes (**Supplementary Table 3**). Consistent with an export/maturation bottleneck, integrative analysis showed that transcripts with strong QKI eCLIP signal often display reduced protein abundance but increased enrichment in the nuclear RNA fraction upon QKI KO (**Fig. 3i**), and sarcomere genes overall were significantly shifted towards increased nuclear expression (**Fig. 3j**). Genes containing cassette exons differentially included upon QKI knockout also showed a modest but statistically significant nuclear enrichment of their host transcripts (**Extended Data Fig. 6d,e**), indicating that splicing failure upon QKI knockout correlates with nuclear retention.

The altered RNA splicing and increased nuclear retention of *TTN*, the longest gene in the human genome and a central architectural scaffold of the cardiac sarcomere, was particularly notable. *TTN* encodes a multi-million Dalton elastic protein that spans half the sarcomere, coordinating sarcomere assembly, stability, and passive elasticity, and mutations in *TTN* are a leading cause of familial cardiomyopathies, including dilated and hypertrophic forms^56^. The extensive QKI binding and multi-exon splicing changes observed in our nuclear RNA-seq of QKI KO (**Fig. 3h**) extend prior findings indicating QKI control of *TTN* splicing^33^ and suggest that *TTN* is dramatically altered upon QKI loss and nuclear-retained. Immunofluorescence staining also revealed near-complete loss of TTN protein along with ACTN2 in QKI-deficient cells (**Fig. 3k**), supporting a model in which defective splicing and nuclear sequestration of *TTN* transcripts underlie the failure to assemble sarcomeres in the absence of QKI. The importance of this QKI-mediated control of *TTN* is further emphasized by recent work^57^, which revealed that nuclear *TTN* pre-mRNA forms an RNA factory that nucleates a variety of RNA processing factors in cardiomyocytes and is characterized by QKI co-localization. Using single-molecule fluorescence in situ hybridization (smFISH) combined with immunofluorescence, we also observed that *TTN* transcripts form discrete nuclear foci that encircle QKI-enriched puncta, consistent with localized accumulation at the *TTN* transcription site (**Extended Data Fig. 6f)**. In contrast, for another QKI-regulated transcript, *RYR2,* we observed distinct, spatially segregated foci that do not colocalize with either *TTN* or QKI (**Extended Data Fig. 6f)**. These findings indicate that QKI–*TTN* condensates may reflect transcript-specific nuclear compartments, rather than generalized hubs for all QKI-dependent transcripts, reflecting a spatial selectivity of QKI–RNA interactions in cardiomyocytes.

Together, these results highlight that the exclusion of exons upon QKI loss not only drives altered isoform expression or decreased RNA abundance likely via nonsense-mediated decay; rather, QKI loss leads to a broad disruption of pre-mRNA processing that often results in nuclear retention. The subsequent failure to produce proteins necessary for sarcomere architecture and cardiac function links QKI-mediated exon inclusion to the sarcomere defects observed in QKI KO cardiomyocytes.

### Intronic QKI binding near 5ʹ splice sites ensures accurate exon inclusion

To define the regulatory rules for QKI-dependent splicing events, we integrated QKI eCLIP-seq binding data with splicing changes in the corresponding differentiation stage to construct cell-type-resolved RNA splice maps^58, 59^. These splicing maps indicated that splicing outcomes were tightly related to QKI binding position: exon inclusion upon QKI KO was characterized by QKI binding near the 5ʹSS (indicating that QKI binding to the 5ʹSS induces inclusion), whereas QKI KO-dependent exclusion showed enriched QKI binding near the 3ʹSS (indicating QKI binding to the 3ʹSS induces exon exclusion) (**Fig. 4a,b**). This positional logic was consistent across all stages of cardiac differentiation and aligned with previously reported motif-predicted patterns from heart^36^ and CLIP-identified maps in neuronal stem cells^60^ and ENCODE-profiled HepG2 cells^52^ (**Fig. 4a**) Notably, intronic QKI eCLIP enrichment flanking the 5ʹSS of exons excluded upon QKI knockout was a consistent pattern of many events in critical cardiac genes, including *ACTN2, NEBL, TTN, RYR2, NIN, CAPZB, TPM1, DST, CASK*, and *CLASP2,* among others (**Fig. 3e,f,h, Fig. 4b, Extended Data Fig. 5a-d, Extended Data Fig. 6a,b**). Matching our results from analysis of all differential splicing events (**Fig. 3d**), genes containing exons excluded upon QKI knockout that had flanking intronic QKI peaks from eCLIP were enriched for cytoskeletal and sarcomeric genes essential for cardiac function (**Fig. 4c**).

**FIGURE 4.**
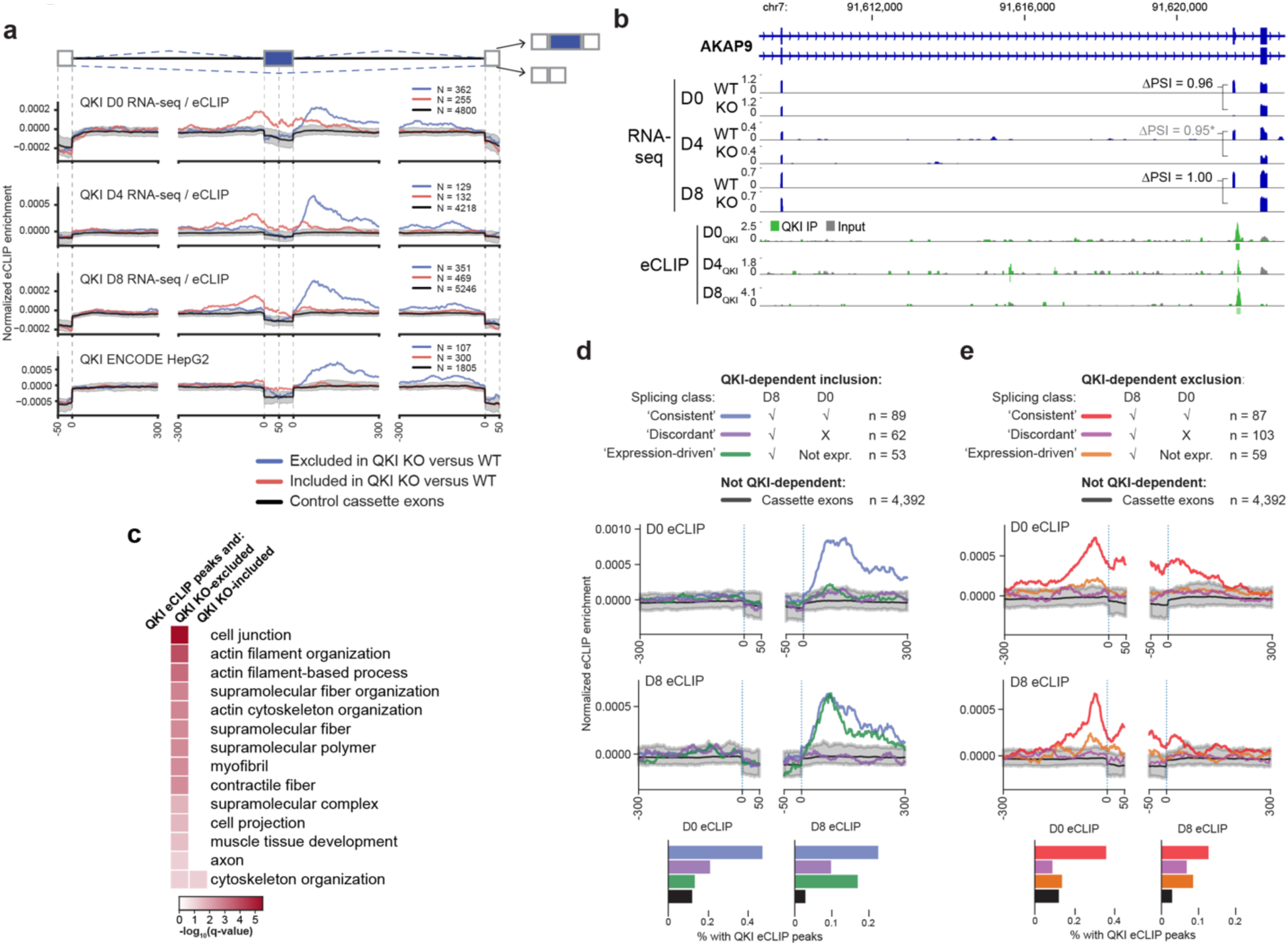
| QKI controls a conserved splicing regulatory program through binding to the 5ʹSS- and 3ʹSS-adjacent intronic regions. **a**, ‘Splicing maps’ for QKI generated by integrating stage-matched eCLIP and splicing data reveal positional logic for QKI-mediated regulation. QKI binding near the 5ʹSS promotes exon inclusion, whereas binding near the 3ʹSS promotes exon exclusion. **b**, Read density of QKI eCLIP and RNA-seq at alternative exons show examples of 5ʹSS-proximal binding at exon 4 of *AKAP9* leading to decreased (QKI-dependent) inclusion upon QKI loss. **c**, Reactome pathway analysis and Gene Ontology (GO) enrichment analysis of QKI-dependent splicing events in cardiomyocytes, stratified by QKI binding position and splicing outcome. Exons bound by QKI near the 5ʹSS and excluded upon QKI loss (QKI-dependent inclusion) are highly enriched for genes involved in sarcomere organization and cardiac contractility. In contrast, exons bound near 3ʹSS and aberrantly included in QKI KO cells (QKI-dependent exclusion) are enriched for neuronal and migration-associated genes. **d,e**, Schematic of classification logic used to assign QKI-bound, dependent (**d**) exon inclusion or (**e**) exon exclusion events into “consistent,” “discordant,” or “expression-driven” categories by comparing splicing and binding at D0 and D8. (center) Splicing maps generated for the indicated categories comparing D0 and D8 indicate that consistent events show consistent QKI binding in D0 and D8, whereas discordant events appear to be indirectly regulated by QKI. D8 exclusive exon inclusion events (green) show enriched QKI binding at D8 but not D0, indicating QKI control of splicing but not underlying gene expression at these events.

As we previously noted that cardiomyocyte differentiation induced a unique QKI interaction landscape (**Fig. 2h**), we next queried the relationship between QKI binding, cell type specificity in transcript expression, and splicing dependency. Comparing the most distinct stages, D0 (pluripotency) and D8 (cardiomyocytes), we found that similar to what we observed for peaks (**Fig. 2h**) there were both groups of “consistent” events (with QKI-responsive splicing throughout differentiation) as well as “expression-driven” events (differentially spliced in D8 but not expressed in D0) (**Fig. 4d,e**). For QKI-dependent exon inclusion events, both classes showed enriched QKI binding in cardiomyocytes (**Fig. 4d**). The expression-driven group contained cardiomyocyte-specific transcripts encoding core sarcomeric components and regulators of contractility, including *ACTN2, CACNA1C, TNNT2,* and *SVIL* (**Fig. 4d**). Thus, there is a unique set of transcripts that are both selectively expressed in cardiomyocytes and rely on QKI for accurate exon inclusion, providing an explanation for the sarcomere disorganization observed upon QKI loss in cardiomyocytes. In contrast, ‘discordant’ exons that were QKI-dependent in D8 but not D0 did not show enriched QKI binding (**Fig. 4d**). Consistent with QKI being expressed in both D0 and D8, this suggests that for the discordant exons, the D8-specific dependence on QKI may occur indirectly through QKI regulation of a different cardiac-specific RBP. QKI-dependent exon exclusion events showed a similar pattern - QKI binding was enriched (proximal to the 3ʹSS) at consistent events but not discordant ones (**Fig. 4e**).

Together, these results reveal a spatial and context-dependent logic for QKI-mediated splicing: binding near 5ʹ splice sites promotes exon inclusion in both a cross-cell type set of QKI targets as well as a specific set of essential cardiac genes (**Fig. 4c,d**), while binding near 3’ splice sites facilitates the exclusion of exons in genes characterized by non-cardiac roles (**Fig. 4c,e**). Despite pervasive QKI binding and widespread presence of its recognition motif across the transcriptome, QKI exerts highly selective control over exon inclusion through proximal intronic engagement at a restricted set of key cardiac transcripts required for cardiac function. This remarkable functional specificity, emerging from widespread binding, points to an unrecognized mechanistic logic in exon recognition—one that requires QKI to convert proximal intronic binding into precise exon inclusion at loci essential for tissue-specific function.

### QKI reinforces U6 pairing at suboptimal 5′ splice sites to drive exon inclusion

We next asked what distinguishes the QKI-dependent exons in essential cardiac transcripts that require QKI for accurate splicing. A central determinant of exon recognition is base-pairing between the 5ʹSS and small nuclear RNAs̶primarily U1, but also U6, which engages a stretch of nucleotides near the beginning of the intron^2, 19, 22, 28^ (**Fig. 5a**). While most 5ʹSS exhibit strong complementarity to U1 and U6, a substantial proportion of human exons harbor intrinsically ʻweak’ 5ʹSS with poor complementarity^29, 61^. Recent work has highlighted evolutionarily conserved weak 5ʹSS in developmentally essential exons̶particularly in heart and brain̶and proposed a regulatory interplay with nearby QKI binding sites, though the experimental validation and mechanism have remained unresolved^6, 29^.

**FIGURE 5.**
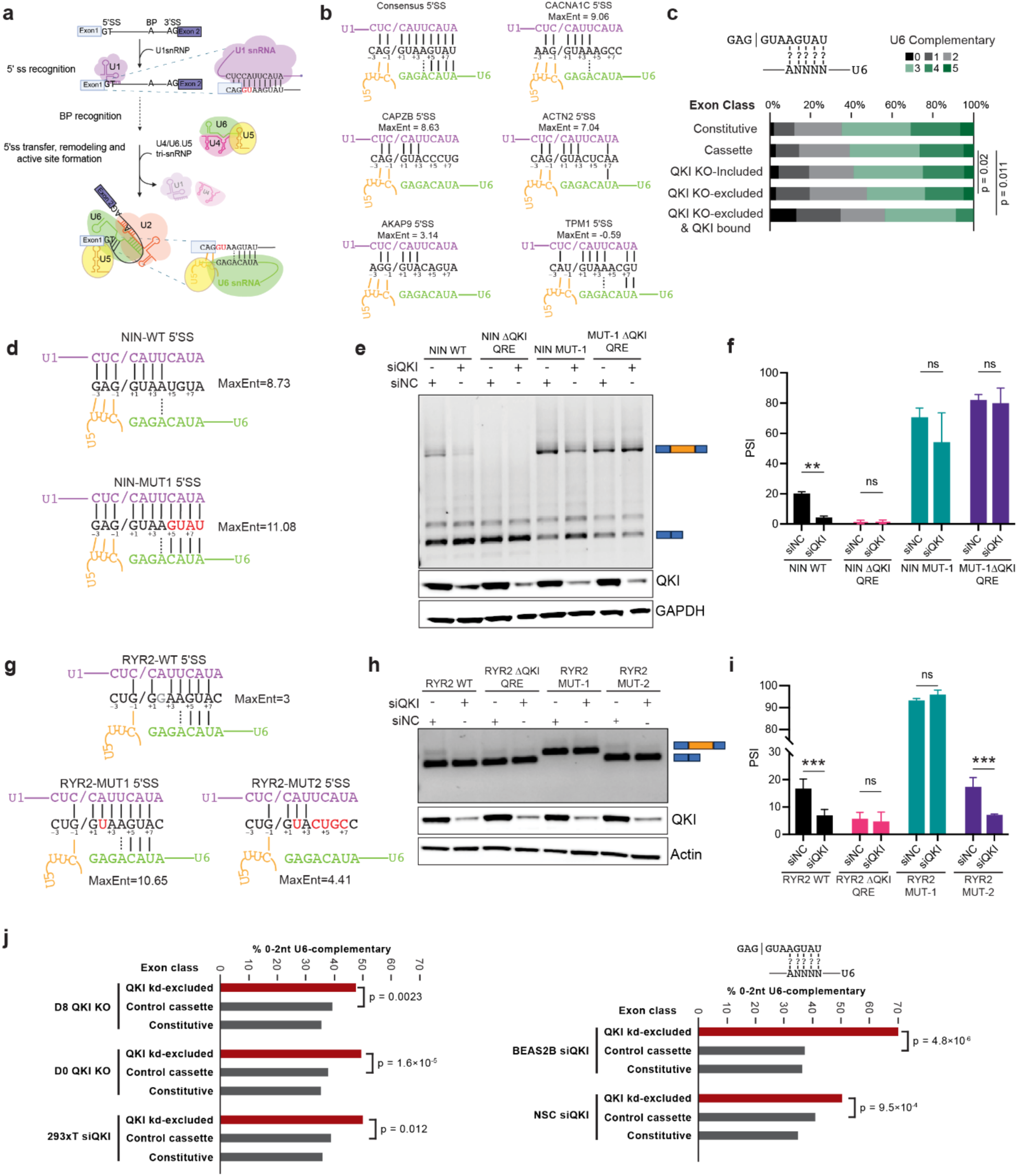
| QKI enhances exon inclusion at suboptimal 5ʹ splice sites with poor U6 complementarity. **a**, Schematic indicating key steps of snRNA recognition of the 5ʹSS, with insets indicating complementarity between canonical 5ʹSS and (top) U1 or (bottom) U6 snRNA. **b**, Illustration of base pairing interactions of U1, U5, and U6 snRNAs at a canonical 5ʹSS and five example exons with QKI-dependent inclusion and QKI eCLIP peaks near the 5ʹSS. Indicated splice site scores are from Maximum Entropy (MaxEnt) modeling. **c**, For indicated exon classes, bars show the fraction of exons with their 5ʹSS containing the indicated number of complementary nucleotides to U6 snRNA. Significance of QKI-regulated exon enrichment for weak U6 base pairing was calculated by Fisher’s Exact Test comparing high (3-5 nt) versus low (0-2 nt) complementarity. **d**, Schematic of wild-type and mutant *NIN* 5ʹSS minigenes, illustrating enhanced U6 base-pairing in MUT1 to test the functional relevance of U6 complementarity in QKI-dependent exon inclusion. Reporters spanning endogenous exons and flanking introns were transfected into HEK293XT cells with QKI siRNA knockdown or non-targeting control. **e**, Representative image of minigene RT-PCRs using wild-type (WT) and U6-optimized (MUT1) *NIN* reporters with or without adjacent QKI-binding motifs, either in the presence or absence of QKI. Bottom, immunoblot confirming QKI knockdown; GAPDH serves as a loading control. **f**, Bar graph representing densitometric quantification of minigene RT-PCRs in (**f**) for *NIN*, n=3. Significance was determined by two-way ANOVA; *p* < 0.01(**), *p* ≥ 0.05 (ns) **g**, Schematic of wild-type (WT), U6-enhanced (MUT1), and U6-weakened (MUT2) *RYR2* 5ʹ splice site minigene constructs, designed to test the functional relevance of U6 base-pairing strength in QKI-dependent exon inclusion. MUT1 improves, and MUT2 reduces, U6 complementarity relative to WT. **h**, Representative image of minigene RT-PCRs using wild-type (WT) and U6-modified (MUT1, MUT2) *RYR2* reporters with enhanced or reduced U6 complementarity, respectively, either in the presence or absence of QKI. Bottom, immunoblot confirming QKI knockdown; Actin serves as a loading control. **i**, Bar graph representing the densitometric quantification of the minigene RT-PCRs in (**f**) for *RYR2*, n=3. Significance was determined by two-way ANOVA; *p* < 0.001(***), *p* ≥ 0.05 (ns). **j**, As in (**c**), predicted U6 complementarity was scored for QKI knockout- or knockdown-excluded exons from our profiling of D8 and D0 QKI knockout cells, our profiling of QKI siRNA knockdown in 293T cells, and published QKI knockdown in BEAS2B and neuronal stem cell (NSC) cells^60, 64^. Across diverse human cell types and datasets, exons excluded upon QKI loss consistently harbor weak 5ʹSS with reduced U6 base-pairing potential, indicating a conserved requirement for QKI in stabilizing U6 engagement at structurally suboptimal 5ʹSS. Significance was determined by Fisher’s Exact Test.

To test whether QKI-dependent cardiac targets preferentially include such weak splice sites, we first systematically analyzed 5ʹSS strength using Maximum Entropy (MaxEnt)^61^ scoring. As expected, cassette exons had lower MaxEnt scores compared to constitutive exons (**Extended Data Fig. 7a**). Notably, exons with QKI-dependent inclusion (ΔPSI ≥ 0.1) showed even weaker 5ʹSS (**Extended Data Fig. 7a**). Restricting the analysis to the fraction of exons that were also flanked by QKI eCLIP peaks showed a less clear pattern, with similar median to cassette exons overall but a subset with particularly weak splice sites (**Extended Data Fig. 7a**).

However, when we inspected individual examples of QKI-dependent cardiac targets, we surprisingly observed that even beyond ‘weak’ 5ʹSS (e.g. *AKAP9, TPM1, CLASP2*), many exons (including examples in *CACNA1C, ACTN2, RBFOX2, CAPZB*, and *SVIL)* with ‘strong’ 5ʹSS by MaxEnt score had strong complementarity to U1 snRNA in the nucleotides immediately adjacent to the splice site but poor complementary to U6 snRNA-interacting nucleotides downstream (**Fig. 5b, Extended Data Fig. 7b**). This also included multiple examples that lack the highly conserved +4A that interacts with the key N-6-modified A-43 on U6^22, 32, 62^. To further dissect the splice site strength in relation to individual snRNAs that interact with the 5ʹSS, we separately modeled U6 and U5 complementarity. For modeling purposes, complementarity to U6 snRNA was defined as the presence of a +4A and complementarity to positions +5 to +8, whereas complementarity to U5 snRNA was modeled based on the last three exonic nucleotides (**Fig. 5c, Extended Data Fig. 7c**), as in previous efforts to separate these classes of 5ʹSS^32, 62, 63^. This confirmed that QKI-bound exons with QKI-dependent inclusion were disproportionately enriched for weak U6 base-pairing potential: while only 35% of all cassette exons had ≤2 predicted U6 base pairs, this proportion rose to 47.6% among QKI-regulated exons and 56.5% among QKI-bound regulated exons (**Fig. 5c**). In contrast, U5 complementarity showed no correlation with QKI dependence (**Extended Data Fig. 7c**). These data support a model in which QKI promotes exon recognition by specifically reinforcing U6 engagement at weak 5ʹSS.

To directly test this hypothesis, we used minigene splicing reporters for two QKI target exons: NIN exon 18 and *RYR2* exon 75. As minigenes recapitulated QKI-dependent inclusion in both 293T and C2C12 (**Extended Data Fig. 5e**), we performed minigene assays in 293T due to improved transfection and knockdown efficiency. Both *NIN* exon 18 and *RYR2* exon 75 are strongly dependent on QKI for inclusion (ΔPSI > 0.9 in cardiomyocytes), yet differ in their snRNA complementarity. *NIN* exon 18 has an approximately median 5ʹ splice site score transcriptome-wide (MaxEnt score of 8.73 versus median 8.45); however, it has good complementarity to U1 but poor complementarity to U6, with only 1 predicted U6-interacting nucleotide (**Fig. 5d**). Similar to previous observations in our cardiomyocyte data, *NIN* exon 18 inclusion in the WT reporter showed a strong (4-fold) dependence on QKI expression (from 20% inclusion in control to 5% inclusion upon QKI knockdown), with complete inclusion loss upon deletion of the QKI motif (**Fig. 5e,f**). In contrast, introducing a 4-nt mutation (TGTA>GTAT) at the 5ʹSS (MUT1) to enhance U6 base-pairing led to enhanced exon inclusion even upon QKI knockdown, reducing QKI dependency from 4-fold to ∼1.5-fold (**Fig. 5e-f**). Furthermore, the enhanced inclusion observed with improved U6 base-pairing in *NIN* MUT1 no longer showed dependence on the QKI binding site (**Fig. 5e,f).** Thus, QKI-dependent inclusion of *NIN* exon 18 can be circumvented by improving U6 complementarity to the 5ʹSS.

The *RYR2* exon 75 5ʹSS contains a non-canonical +2G (GG) instead of the typical GT in the first two intronic nucleotides and is thus predicted to have weak U1 engagement but relatively stronger U6 pairing (4 potential base pairs) (**Fig. 5g**). In HEK293 cells, the WT *RYR2* exon 75 reporter showed QKI-dependent inclusion as expected (**Fig. 5h,i**). Conversion of the 5ʹSS +2G to the more canonical +2T (*RYR2* MUT1) or +2C (*RYR2* MUT3) resulted in consistently high inclusion levels that were not dependent on QKI (**Fig. 5h,i**, **Extended Data Fig. 7d,e**). However, a second mutation (*RYR2* MUT2), which preserved the +2T / U1 complementarity but disrupted other U6-interacting bases, restored QKI dependence and reduced inclusion (**Fig. 5h,i**). Thus, similar to *NIN*, U6 complementarity (rather than simply MaxEnt score or U1 complementarity) was the critical determinant of QKI dependence at *RYR2* exon 75. In contrast, mutations to the *NIN* exon 18 minigene, which increased U6 complementarity, were not able to rescue the splicing defect observed with mutation of the highly conserved +2T nucleotide, highlighting context-specificity for the ability of QKI to rescue poor U1 complementarity **(Extended Data Fig. 7f-i)**.

Our minigene studies indicated that the U6-dependent splicing logic for QKI observed in cardiomyocytes extended to HEK293T cells. To test whether this extends to other cellular contexts where QKI plays critical physiological roles, we performed eCLIP and siRNA knockdown of QKI, followed by RNA-seq in 293T cells, coupled with systematic reanalysis of multiple publicly available datasets that combine QKI binding (eCLIP) with QKI perturbation (knockout or knockdown) and splicing readouts^60, 64^. Across diverse cellular contexts, exons excluded upon QKI loss consistently harbored 5′SS with weaker U6 base-pairing potential, mirroring our observations in cardiomyocytes (**Fig. 5j**). Notably, this dependency on U6 snRNA base-pairing potential was not observed for U5 base-pairing in the same datasets, reinforcing the model that QKI selectively stabilizes exon inclusion specifically at U6-sensitive exons (**Extended Data Fig. 7j**). These results reveal a universal molecular logic for QKI in promoting splicing fidelity by facilitating U6 engagement at suboptimal 5′ splice sites and establish QKI as a broadly acting regulator of exon inclusion across cell types. Together, these findings uncover a mechanistic link for QKI-mediated control of exon inclusion in the heart, whereby QKI selectively binds near 5′SS with poor U6 complementarity to enable snRNA engagement and ensure splicing fidelity for exons essential for sarcomere assembly and contractile function.

### QKI directly engages U6 snRNA independently of its intronic binding activity

Splice site recognition at a weak 5ʹSS depends on coordinated engagement by both U1 and U6 snRNAs, with U6 playing a catalytic role as part of the tri-snRNP complex. Given that QKI-dependent exons were specifically sensitive to U6 complementarity, we hypothesized that QKI might physically interact with U6 snRNA to stabilize splice site recognition during cardiac differentiation. To assess whether QKI directly engages U6 snRNA during human cardiac specification, we systematically reanalyzed our eCLIP datasets across differentiation stages. This analysis revealed robust enrichment of reads mapping to U6 snRNA at all time points (**Fig. 6a**), with negligible enrichment of other spliceosomal snRNAs (**Fig. 6b**), indicating a selective and stable interaction between QKI and U6. To benchmark the enrichment and specificity of this interaction, we compared the enrichment of U6 in QKI eCLIP to the 155 ENCODE-profiled RBPs^50, 52^. This analysis revealed that QKI was one of the most highly-enriched RBPs, with enrichment comparable to that of SMNDC1 (**Fig. 6c**), a well-characterized splicing factor associated with the tri-snRNP^65^. In addition, reinforcing the specificity of interaction of QKI with U6, we found that QKI does not associate with its minor spliceosome counterpart U6ATAC, in contrast to proteins such as SMNDC1 that bind to both snRNAs (**Fig. 6d**). Interestingly, QKI binding to U6 was consistently enriched near the 5ʹ end of U6 snRNA across all stages of cardiomyocyte differentiation as well in K562 and HepG2 cells profiled in ENCODE (**Fig. 6e**). The 5ʹ end of U6 snRNA harbors a conserved stem-loop structure and is adjacent to the 5ʹSS interacting site, consistent with this interaction having functional relevance to splice site recognition (**Fig. 6e**). This contrasts with other tri-snRNP-associated splicing factors such as PRPF8, RBM22, SMNDC1, and BUD13, which are predominantly enriched at the 3’ region of U6 snRNA (**Fig. 6e**). These findings position QKI as a U6-associating RBP with binding and specificity on par with dedicated components of the U6 snRNP, but with a distinct positional signature of QKI‒U6 interaction from these factors.

**FIGURE 6.**
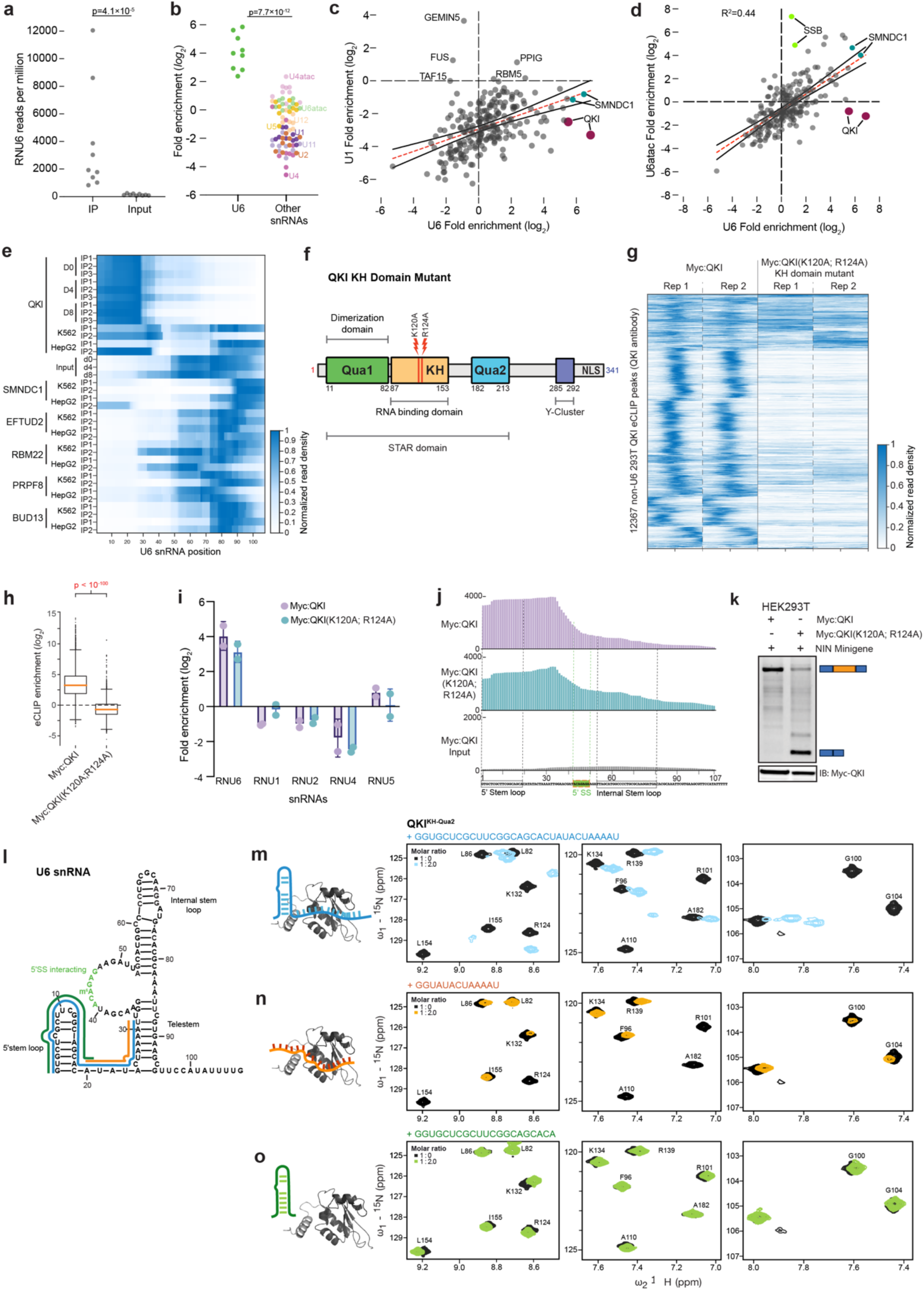
| QKI directly engages U6 snRNA independent of Its intronic binding activity. **a**, QKI eCLIP-seq reveals robust and selective binding to U6 snRNA across all stages of human cardiac differentiation (D0, D4, D8). Significance was determined by the Mann-Whitney *U* test. **b**, QKI shows negligible enrichment for other spliceosomal snRNAs (U1, U2, U4, U5, U6ATAC, U11, U12, U4atac) compared to U6 snRNA across all stages of human cardiac differentiation. Significance was determined by the Mann-Whitney *U* test. **c,** Comparative enrichment analysis of QKI on U6 snRNA (log₂ fold change over size-matched input) relative to 155 ENCODE-profiled RBPs, showing selective U6 engagement (comparable to canonical U6-binding factor SMNDC1) but not to U1 **d**, QKI shows selective binding to U6 snRNA, but not to the minor spliceosomal U6ATAC snRNA, in contrast to SMNDC1, which binds to both snRNAs. **e**, QKI binds specifically to the 5ʹ region of U6 snRNA across all stages and replicates, distinct from canonical U6-binding splicing factors (e.g., PRPF8, RBM22), which predominantly bind the 3ʹ region. **f**, Domain architecture of QKI and schematic of the KH domain double mutant (K120A/R124A) used to assess functional contributions of canonical RNA binding to U6 engagement. **g**, Normalized read density in eCLIP of transgenic MYC-tagged wild-type and KH domain mutant QKI in HEK293T cells, shown for eCLIP peaks identified in independent experiments with anti-QKI antibody in HEK293T. **h**, Quantification of (**g**) shows a significant loss of eCLIP signal with the KH domain mutant. **i**, Enrichment of QKI binding at indicated snRNAs in cells expressing WT or KH mutant QKI indicates that U6 binding is preserved upon mutation of the canonical RNA recognition domain. **j**, Base-resolution eCLIP mapping of QKI binding to U6 snRNA reveals that the selective enrichment at the 5ʹ stem-loop and adjacent single-stranded region critical for 5ʹSS engagement is preserved in the QKI KH domain mutant. **k**, RT-PCR using a *NIN* exon 18 minigene shows that the KH-domain mutant QKI fails to restore splicing despite retaining U6 binding. **l,** Schematic showing secondary structure of human U6 snRNA with key regions including the 5ʹ stem-loop, 5ʹSS interacting domain, internal stem-loop, and telestem. Colors indicate regions used for NMR analysis. **m-o**, NMR titration of recombinant QKI KH–QUA2 protein with *in vitro* transcribed U6 RNA fragments. Residues affected based on chemical shift perturbations overlap with the canonical RNA-binding surfaces. **m**, The U6 fragment containing both the 13-mer sequence with a partial QRE motif and the 5ʹ stem-loop induces the strongest chemical shift changes, consistent with an extended RNA-binding interface and a higher affinity interaction. **n**, The 13-mer alone also triggers pronounced shifts, yet to a lesser extent compared to (**m**). **o**, In contrast, the 5ʹ stem-loop alone causes only minor perturbations, consistent with a weak interaction. Error bars represent ±SEM; *p-*values are calculated using Student’s *t*-test; biological replicates *n* = 3

QKI has previously been shown to bind RNA primarily via its canonical RNA binding KH domain, which interacts with an intronic QKI response element (QRE) that is often found as a bi-partite motif bound by dimerized QKI^47^. To determine whether QKI interaction with U6 utilized the same mechanism, we over-expressed either WT myc-tagged QKI-5 or a myc-QKI-5 (K120A; R124A) KH domain mutant previously shown to disrupt RNA recognition via the KH domain^66^ and performed eCLIP-seq in HEK293T cells (**Fig. 6f**). As expected, WT QKI showed robust binding to intronic regions bound in eCLIP of endogenous QKI, while the KH mutant exhibited a marked loss of intron binding at these sites (**Fig. 6g, 6h**), confirming disruption of canonical intronic RNA interaction. Remarkably, however, U6 snRNA binding persisted in the QKI KH domain mutant (**Fig. 6i**), indicating that QKI-U6 engagement occurs independently of intronic RNA-binding activity. Mapping the binding of QKI to U6 at base pair resolution again indicated that both QKI WT and KH domain-mutant bind specifically to the 5ʹ region of U6 snRNA, adjacent to but not overlapping the 5ʹ splice site recognition sequence (**Fig. 6j)**. Given that the QKI KH domain mutant maintains U6 interaction, failure to induce exon inclusion of *NIN* exon 18 (**Fig. 6k**) confirms that QKI function depends on dual engagement of both U6 and the pre-mRNA QRE. As the QKI KH domain is the primary RNA-binding module, it is plausible that in a dimerized context, QKI recognizes intronic RNA and U6 snRNA through overlapping, yet adaptable interfaces that utilize the same QKI domain.

To confirm the QKI–U6 RNA interaction and map the binding interface of QKI with U6 snRNA, we performed Nuclear Magnetic Resonance (NMR) titrations. For this, we monitored spectral changes of amides in recombinant ^15^N-labeled QKI KH–QUA2 with *in vitro* transcribed fragments of the U6 RNA that correspond to key regions in U6 snRNA (**Fig. 6l**). Upon stepwise titration of U6 RNA, we observed chemical shift changes indicating a specific interaction (**Extended Data Fig. 8a**). Using a large fragment of U6 that included both the 5ʹ stem-loop and neighboring linear sequence region, we observed spectral changes for residues in the canonical RNA-binding surface of the KH domain, including the a1 helix, GXXG loop, a2 helix, and the QUA2 helix (**Fig. 6m**, **Extended Data Fig. 8a)**. A U6 fragment containing only the single-strand RNA linear sequence adjacent to the hairpin (which also harbors an ACUAAA sequence that resembles the core ACUAAY QKI motif) also induced significant spectral changes (**Fig. 6n**, **Extended Data Fig. 8a)**. The enhancement of these changes in the presence of the U6 5ʹ stem-loop (**Fig. 6m**) presumably reflects additional stabilizing contacts. However, a fragment containing only the U6 5ʹ stem-loop (U6 SLI RNA) induces only minor chemical shift changes, indicative of a relatively weak interaction (**Fig. 6o**, **Extended Data Fig. 8a; last panels**). NMR spectral changes are also observed for the amide of R124 located in the b2 strand of the KH domain, a residue previously implicated in canonical RNA binding^66^. However, as the mutation of this residue to alanine in the K120A/R124A mutant did not impair U6 binding in eCLIP assays (**Fig. 6i,j**), these results suggest that this residue alone may not be critical for U6 interaction in this context. Together, these data reveal an RNA-binding interface involving QKI KH-QUA2 with the 13-mer hairpin-adjacent linear region in U6, which is further enhanced by the 5ʹ proximal U6 RNA stem loop. This demonstrates a direct interaction between QKI and U6, which is functionally and spatially distinct from its recognition of intronic QRE motifs.

As QKI has previously been shown to act via dimerization and binding to a bipartite sequence composed of a combination of two neighboring QKI motifs^66^, this suggests that the QKI interaction might bridge U6 snRNA with an intronic QRE to promote splicing. In silico modeling of molecular docking further supports this mode of interaction, positioning U6 within a positively charged groove formed by the QKI dimer, with the 5ʹ region situated adjacent to NMR-identified residues (**Extended Data Fig. 8b**). Using this, we modeled how U6 snRNA could fit within the three-dimensional surface of the QKI dimer. By simulating spatial complementarity and electrostatic compatibility, the analysis revealed that U6 can dock into a positively charged groove on QKI with its 5′ region aligning along residues identified by NMR as undergoing chemical shifts upon RNA binding (**Extended Data Fig. 8c**). This in silico reconstruction provides a mechanistic framework for how QKI can interact with U6, revealing a unique mode of RNA recognition involving the 5ʹ end of U6 which appears to be distinct from spliceosome complex components such as SMNDC1, RBM22, and PRPF8, which instead show enriched binding at the 3′ end of U6. Instead, QKI binds the 5ʹ stem-loop region proximal to the splice site recognition domain, ideally placed to stabilize U6 pairing with pre-mRNA.

Together, these findings support a model in which QKI acts as a molecular bridge, simultaneously recognizing intronic motifs and U6 snRNA via its KH–QUA2 domain to stabilize early spliceosome assembly at weak 5ʹ splice sites that lack strong complementarity to U6 snRNA. This dual interaction suggests a coordinated mechanism in which QKI dimers may first bind intronic QREs and subsequently engage U6 snRNA as the tri-snRNP complex arrives—providing a temporally dynamic scaffold that reinforces splice site pairing and splicing fidelity.

### Native complexome and IP–MS map QKI interactions with the pre-catalytic spliceosome

The formation of a stable U6:5ʹSS duplex acts as a molecular checkpoint for catalytic activation^67,68^. Given our observations that QKI directly interacts with U6 and enhances splicing at suboptimal 5ʹSSs with weak U6 pairing, we hypothesized that along with this QKI might directly associate with tri-snRNP components and early spliceosomal factors to enable tri-snRNP progression.

To assess whether QKI physically associates with spliceosomal components in an unbiased manner, we first employed an RNA-aware nuclear complexome approach (**Fig. 7a**). Nuclei were isolated from UV-crosslinked WT and QKI knockout (KO) cells under physiological buffer conditions, and native protein complexes were resolved by BN-PAGE and analyzed by mass spectrometry, with and without RNase treatment. This allowed us to resolve the native protein complexes that associate with QKI and to distinguish between RNA-dependent and independent co-associations.

**FIGURE 7.**
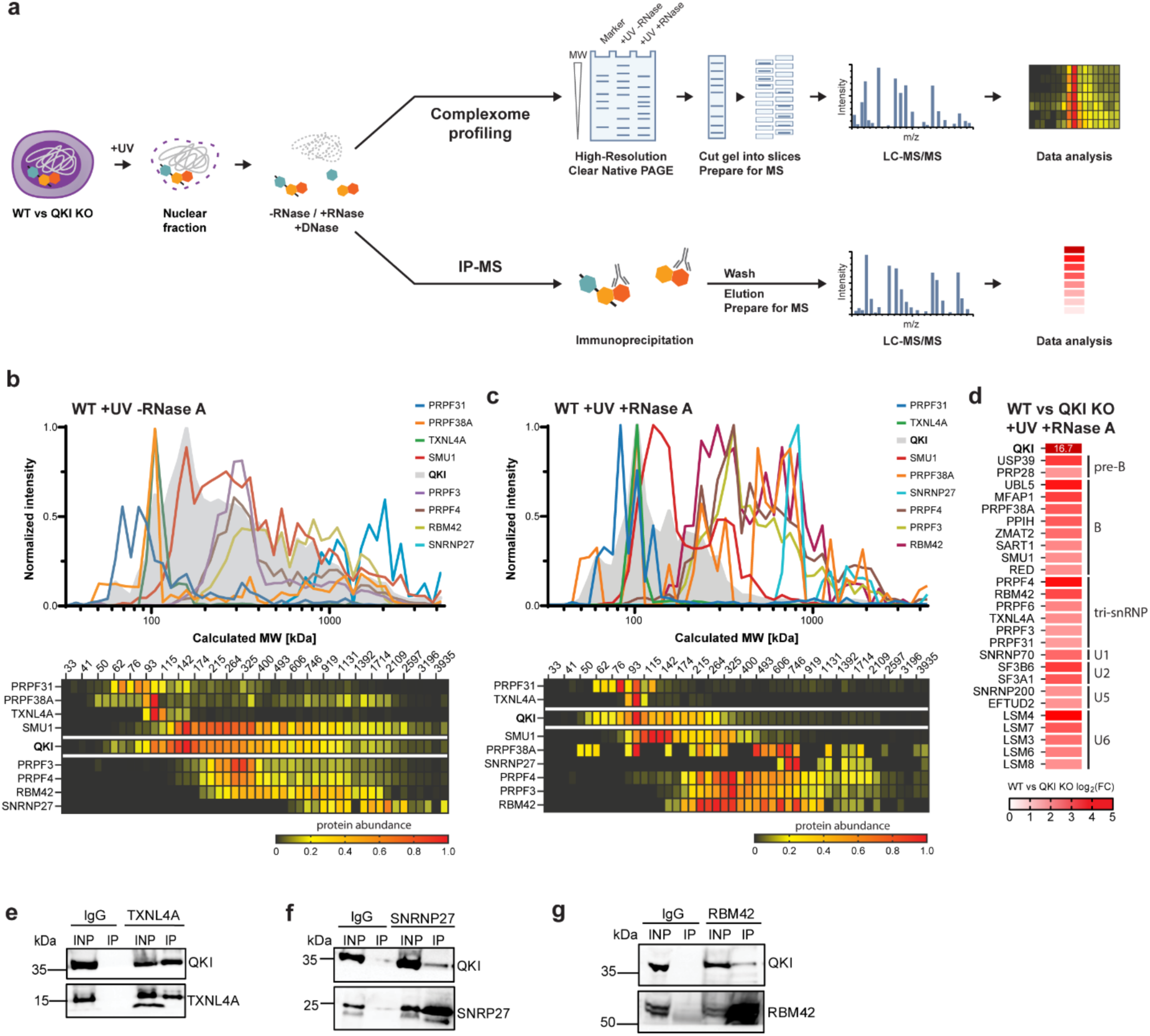
| QKI integrates into native spliceosomal complexes and enforces U6 engagement at weak 5ʹ splice sites. **a**, Schematic of the RNA-aware nuclear complexome and IP-proteome workflows. UV-crosslinked nuclei from WT and QKI KO cardiomyocytes were subjected to BN-PAGE and LC-MS/MS with or without RNase treatment (complexome), or immunoprecipitation of endogenous QKI followed by RNase-treated LC-MS/MS (IP proteome). **b**, RNA-aware complexome profiling without RNase reveals extensive co-migration of QKI with spliceosomal assemblies, including PRPF38A, TXNL4A, USP39, SMU1, RBM42, and SNRNP27, suggesting RNA-mediated interactions with multiple splicing stages. **c**, RNase-treated complexome narrows the QKI interactome, retaining defined associations with early spliceosomal and tri-snRNP proteins, indicating direct protein-protein interactions independent of RNA. **d**, RNase-treated immunoprecipitation proteome confirms a coherent subset of QKI interactors— TXNL4A, PRPF4, SMU1, RBM42, USP39, and SART1, overlapping with complexome results and highlighting QKI association with early spliceosome complexes. **e-g**, Western blot validation of QKI co-immunoprecipitation with (**e**)TXNL4A, (**f**) SNRNP27, and (**g**) RBM42 confirms direct interaction with QKI.

In the absence of RNase, QKI co-migrated with a wide array of spliceosome complexes, suggesting extensive RNA-mediated interactions (**Fig. 7b**). Notably, QKI co-resolved with key components of the pre-catalytic spliceosome, specifically PRPF38A, PRPF4, TXNL4A, SMU1, SNRNP27, and RBM42^68, 69^—implicating QKI in multiple stages of spliceosome assembly and remodeling (**Fig. 7b**). Upon RNase treatment, the overall interactome narrowed considerably, yet QKI retained co-migration with a defined subset of splicing regulators, indicative of RNA-independent associations likely mediated by direct protein–protein contacts (**Fig. 7c**). These QKI-associated retained complexes included factors critical for U4/U6.U5 tri-snRNP stability (TXNL4A, PRPF4)^69, 70^, 5′SS selection fidelity (RBM42, SMU1)^68, 71^, and B complex activation (PRPF38A)^72, 73^, aligning with our model that QKI acts at the interface of splice site recognition and pre-catalytic activation (**Fig. 7c**).

To define direct interactors with higher confidence, we performed immunoprecipitation (IP) of endogenous QKI from wild type UV-crosslinked nuclei followed by mass spectrometry under stringent RNase treatment in addition to QKI-KO as a control (**Fig. 7a, bottom half**). This IP proteome revealed a reproducible and coherent set of direct QKI interactors, including TXNL4A, PRPF4, SMU1, and RBM42—overlapping significantly with the complexome dataset and reinforcing the association of QKI with pre-catalytic spliceosomal assemblies (**Fig. 7d**). Notably, QKI interactors also included USP39^74, 75^, UBL5^76^, SART1^75^, and SMU1^71^ (**Fig. 7d**), factors involved in recruitment and stabilization of the tri-snRNP complex during handover of the 5′SS from initial recognition to tri-snRNP engagement and B complex formation^69, 73^. These associations support a model in which QKI acts as a regulatory scaffold that secures early 5′SS recognition and promotes productive tri-snRNP docking at suboptimal splice sites.

To independently validate these interactions, we performed co-immunoprecipitation to confirm whether QKI associated with specific tri-snRNP components that were identified in complexome profiling and are present at the interface of 5ʹSS transfer during transition from pre-B to B spliceosomal complexes^68, 69^. These assays confirmed direct interactions between QKI and TXNL4A, SNRNP27, and RBM42 (**Fig. 7e-g**). Together, these findings reveal that QKI can associate with native spliceosomal complexes, interacting both in an RNA-dependent and independent manner with core U6 snRNP and tri-snRNP factors that mediate U6 stability, tri-snRNP integrity, and 5′SS recognition. Their direct association with QKI suggests a mechanistic framework wherein QKI may bridge U6 pairing and spliceosomal integration, acting as a scaffold or adaptor that ensures successful splice site handover and catalytic activation—particularly at weak 5′SS.

Thus, these findings demonstrate that splicing fidelity at suboptimal splice sites is not only an intrinsic property of the spliceosome but can be a developmentally regulated outcome shaped by context-aware auxiliary factors that can directly interface with spliceosome components. By coupling cis-element recognition to U6 engagement, QKI exemplifies how RNA-binding proteins can decode local RNA features to guide core splicing machinery and safeguard transcript integrity across developmental and cellular contexts.

## DISCUSSION

Precise exon recognition is fundamental to splicing fidelity and accurate gene expression. Although the spliceosome is pan-cellular, splicing decisions vary across cell types and developmental stages. Many exons essential for tissue function—particularly in the heart—are alternatively spliced^6^, and alternatively spliced exons are often flanked by weak 5ʹ splice sites that deviate from the consensus motif and would be expected to be poorly recognized^29–31^. Nonetheless, these suboptimal splice sites are reliably recognized and utilized in vivo, pointing to context-specific mechanisms that actively enforce splicing fidelity. Recent studies have shown that even core spliceosomal components undergo dynamic remodeling during development, challenging the view that fidelity is intrinsic to the spliceosome^7, 22, 77, 78^. However, how weak splice sites are accurately recognized, particularly to ensure proper splicing of genes essential during organogenesis, has remained unresolved^29, 79^.

Our study directly addresses this gap by identifying the RNA-binding protein QKI as a molecular scaffold that interacts with both intronic RNA and U6 snRNA, thereby enabling exon inclusion at weak splice sites with poor U6 complementarity and directly linking tri-snRNP recruitment and stability to the presence of intronic cis-regulatory elements. Thus, QKI represents a context-adapted strategy by which splicing fidelity is actively maintained in the developing heart. These QKI-dependent exons are embedded in key sarcomeric genes—including *TTN*, *NEBL*, and *RYR2*— making accurate splicing essential for sarcomere architecture and contractility, in agreement with studies examining QKI function in the mouse heart^34, 36^. QKI, despite being ubiquitously expressed, orchestrates context-dependent functions from embryogenesis to tissue homeostasis^34, 80, 81^. QKI exerts post-transcriptional control over splicing, translation, RNA localization, and mRNA stability, with selectivity partly conferred by distinct QKI isoforms^34–36, 38, 39, 41, 48^. For example, it governs splicing in cardiomyocytes and endothelium, directs RNA localization and myelination in glia and oligodendrocytes, and reprograms splicing during monocyte–macrophage differentiation, placing QKI as a key regulator of embryogenesis and homeostasis^34–36, 38, 39, 41, 48^. Recent studies linked QKI to splicing regulation in the developing mouse heart^34, 35^ and differentiated cardiomyocytes^33, 36^. How QKI simultaneously serves as a general hub of RNA metabolism while also executing highly specific functions, such as controlling cardiac function and contractility, remained unresolved^33–36^. In particular, the lack of transcriptome-wide, time-resolved experimental maps of QKI-RNA engagement across cardiac development has made it difficult to distinguish housekeeping functions from context-specific rules that govern target selection and splicing regulation, ensuring sarcomere architecture and cardiac function

By monitoring gene expression dynamics and developmental phenotyping along a cardiac differentiation time course using QKI KO hESCs together with temporally controlled cardiomyocyte-specific rescue, we show that QKI is dispensable for cardiomyocyte identity but essential for contractile function. QKI-deficient cells robustly activate cardiac transcriptional programs yet fail to activate the cardiac splicing program and assemble sarcomeres, revealing a mechanistic bottleneck at weak 5ʹ splice sites that disrupts functional maturation. Our stage-resolved eCLIP-seq analysis reveals that the QKI RNA interactome is dynamically remodeled during cardiac differentiation, progressively targeting transcripts essential for sarcomere assembly and contractile function in maturing cardiomyocytes. Rather than generally regulating developmental transitions, QKI enhances splicing fidelity at weak splice sites in a core set of transcripts required for contractility, operating as a molecular bridge between 5ʹSS, intronic recognition elements, and the catalytic core of the spliceosome. Furthermore, we observe that these interactions not only lead to isoform switching but also to nuclear retention and loss of protein products for key sarcomeric proteins, including *TTN*, *NEBL*, and *TNNT2*. Thus, the role of QKI in maintaining accurate splice site recognition at key genes has an impact throughout the life cycle of these RNAs.

Using this data, we uncover a previously unrecognized mechanism by which QKI safeguards splicing fidelity through direct engagement with not only intronic sequence, but also U6 snRNA and the U4/U6.U5 tri-snRNP complex. Previous analysis of the role of QKI in splicing proposed a context-dependent regulatory concept in which QKI binding upstream of an exon leads to repression of the exon, whereas binding downstream results in inclusion^6, 35, 36^. Indeed, mechanisms for RBP-mediated activation of exon inclusion are generally poorly understood^4, 22, 28, 30, 32, 52, 69, 79, 82–84^, with specific models only shown for U1 recruitment by FUS and RBFOX1 and U2 by RBM39^82, 83, 85, 86^.

Our data position QKI as the first alternative splicing regulator to directly reinforce the U6 snRNP, suggesting it acts as a molecular scaffold that reinforces exon recognition by supporting splice site handover to the tri-snRNP. Notably, QKI interacts through both RNA-dependent and RNA-independent contacts not only with U6 itself but also with several tri-snRNP components that sit at the interface between U6 and the 5ʹSS (including RBM42, SMU1, and TXNL4A), suggesting it may play an active role in tri-snRNP remodeling and catalytic commitment. Recent work has described two unique classes of 5ʹ splice sites defined by the strength of interaction with either U5 or U6 snRNA^32, 62, 87^, suggesting that poor U6 complementarity could serve as a novel mechanism through which a tissue-specific splicing regulatory program can be controlled by an individual RBP that can bridge these poor U6:5ʹSS interactions. We identify QKI as a context-aware splice-fidelity factor that secures accurate 5ʹ splice-site choice by directly engaging U6 and interfacing with U4/U6·U5 tri-snRNP components and fidelity modulators (SMU1, USP39, RED, SNRNP27, TXNL4A, RBM42^68, 69, 71, 72, 75, 88, 89^), thereby providing in vivo mechanistic support for the U5/U6 two-class model and providing an explanation for how U6-weak exons can be precisely spliced during cardiac development.

The essential role of accurate splicing in sarcomeric genes is underscored by their frequent disruption in cardiac disease^9, 90^. Genes such as *TTN, RYR2*, and *NEBL* are structurally complex, essential for contractility, and commonly affected by pathogenic splicing variants^90–94^. While mis-splicing of these transcripts has been linked to cardiomyopathies, the regulatory principles that ensure their precise processing during development have remained unclear^15^. Our findings identify QKI as a cardiac contractility-specific fidelity factor that safeguards exon recognition at suboptimal splice sites. These findings align with and build upon recent insights revealing that the fidelity of splicing is not an intrinsic property of the spliceosome, but rather a regulated feature shaped by cellular context^6, 22, 30, 32, 62^. Studies in pluripotent stem cells and differentiating tissues have demonstrated that core spliceosome components, including tri-snRNPs and associated helicases, undergo dynamic remodeling to accommodate tissue-specific splicing demands^72, 78^. By coupling intronic motif recognition to U6 snRNA engagement, QKI ensures accurate splicing of long, intron-rich cardiac transcripts whose misprocessing compromises tissue function.

Building on this context-specific role in the heart, the broader functional versatility of QKI highlights its capacity to program distinct layers of RNA regulation in different cellular environments. These diverse activities—spanning transcript stabilization, translational control, and splice site fidelity— underscore a unifying principle: QKI operates as a context-sensitive recognition module that couples RNA elements to specific downstream processing machineries. Whether engaging the spliceosome at suboptimal 5′ splice sites in cardiomyocytes or regulating mRNA fate in neurons and stressed cells, QKI exemplifies how ubiquitously expressed RBPs can execute highly specialized, spatially localized functions. Our findings reveal that QKI decodes local RNA cues and docks the appropriate RNA-processing machinery, converting cellular context into precise exon decisions during development. In doing so, this work fills a fundamental gap in our understanding of how exon recognition is safeguarded at weak splice sites—particularly in the context of organogenesis—and reframes splicing fidelity as a regulated, context-dependent process. More broadly, our study provides a mechanistic framework for how RNA-binding proteins cooperate with the core splicing machinery in a context-aware manner to ensure splicing fidelity and preserve the integrity of developmental gene expression programs, with implications for understanding congenital diseases, post-transcriptional regulation, and tissue-specific vulnerability to splicing defects.

## Acknowledgements

We thank Dr. Alessandro Annibaldi (CMMC, University of Cologne), Prof. Christoph Dieterich (University of Heidelberg), and Prof. Argyris Papantonis (University of Göttingen) for critical input and discussions throughout this project. We would like to thank Prof. Zefeng Wang (CAS Center for Excellence in Molecular Cell Science, Shanghai Institute for Biological Sciences, Chinese Academy of Sciences for generously sharing splicing reporter constructs and Dr. Sam Fagg (University of Texas Medical Branch) for QKI plasmids. We are grateful to all members of the Van Nostrand and Kurian laboratories for their support, insight, and feedback. This project represents a joint effort and equal contribution between the Kurian and Van Nostrand laboratories. We want to thank Dr. Jan-Wilm Lackmann (CECAD/CMMC Proteomics core) for advice and suggestions on proteomics experiments, and the Cologne Center for Genomics (for assistance with RNA-seq) and CECAD/CMMC Proteomics Facility (for QKI KO proteomics).

## Ethical permissions

Ethical permission to work with human embryonic stem cells for this project has been granted to the Kurian lab by the Robert Koch Institute under AZ 3.04.02/0145.

## Author contributions

Conceptualization: L.K. and E.L.V.N. Methodology: W.Y., M.V.A., D.B., C.Y., J.Z., L.H., C.A., L.R., A.C.S., F.Y., M.D.B., F.Z., H.H., R.S., B.C., J.B., R.B., G.D., A.R., I.W., C.M., M.S., E.L.V.N., and L.K. Investigation: M.V.A., W.Y., E.L.V.N., and L.K. Visualization: E.L.V.N., L.K, M.V.A., and J.Z., Supervision: L.K. and E.V.N, Writing—original draft: L.K., M.V.A, and E.V.N, Writing—review and editing: L.K., E.L.V.N., R.B, M.V.A., and J.Z.

## Data availability

RNA-seq and eCLIP data has been deposited at the Gene Expression Omnibus (GSE306691).

## METHODS

### Generation of CRISPR/Cas9-mediated QKI Knockouts in hESCs

To create CRISPR-Cas9-mediated knockout hESCs, 175,000 cells were seeded per well in a 6-well plate and incubated overnight. The following day, a transfection mixture was prepared, consisting of 200 µl OptiMEM (Thermo Fisher Scientific, 31985-047), 12 µl FuGene HD (Promega, E2311), and 3 µg gRNA plasmid. This mixture was vortexed briefly and incubated at room temperature for 15 minutes. During this incubation, the culture medium was replaced with 2 ml of fresh FTDA growth medium^95^. After incubation, the transfection mixture was added dropwise to the cells, followed by a 24-hour incubation.

GFP expression was then assessed using a cell culture microscope. If at least 10% of the cells were found to be GFP-positive, selection was performed for 24 hours by adding 0.5 µg/ml puromycin to the FTDA growth medium. Following selection, puromycin was removed, and the cells were allowed to recover for 2-3 days.

For the isolation of single-cell-derived QKI knockout clones, clonal dilution was performed, with 2000-6000 cells seeded per well in a 6-well plate containing 1.5 ml FTDA growth medium supplemented with 10 µM Y-27632. After 24 hours, an additional 0.5 ml of this medium was added, and 1 ml was replaced with fresh FTDA the following day. From that point onward, all medium was replaced daily with 1-1.5 ml FTDA until colonies were sufficiently large for screening.

For screening, approximately 20% of each colony was sampled, preferably from the center, and lysed in 30 µl of Quick Extract DNA Extraction solution (Biozym, 101094). The cells were vortexed for 15 seconds, incubated at 65°C for 6 minutes, vortexed again for 15 seconds, and then incubated at 98°C for 2 minutes. For screening PCRs, 5 µl of the resulting genomic DNA was used, while the remaining genomic DNA was stored at -20°C.

Positive clones were allowed to grow further until an appropriate size was reached for picking. To pick colonies, surrounding cells were removed, and the colony was divided into small pieces using a 27-G syringe. Colony pieces were transferred to a fresh well containing 2 ml FTDA growth medium with 10 µM Y-27632. After one week, the colonies were further divided into small pieces and dissociated using Dispase (Invitrogen, 17105-041). The cells were washed once with DPBS, then 1 ml of Dispase was added for a 5-minute incubation at 37°C. After incubation, the colonies were carefully washed twice with DPBS, and 2 ml of FTDA growth medium was added. The cells were scraped from the plate using a cell scraper. The colonies were briefly centrifuged, the supernatant was removed, and the cells were seeded onto a fresh 6-well plate. Once the well was at least 50% covered with colonies, the cells were returned to monolayer culture by splitting with Accutase.

### Cardiomyocyte differentiation

Cardiomyocyte differentiation was conducted following a previously established protocol^95, 96^. In summary, confluent human embryonic stem cells (Hues6 hESCs) were enzymatically dissociated into single cells using Accutase and incubated at 37°C for 10 minutes. The enzymatic reaction was stopped by adding two volumes of F12/DMEM. Cell density was determined using a Neubauer counting chamber, and 230,000 cells per square centimeter (equivalent to 450,000 cells per well in a 24-well plate) were centrifuged at 300 x g for 2 minutes at room temperature.

For D0 (pluripotency), cells were plated in FTDA growth medium^95^, and cultured overnight prior to UV crosslinking. For cardiomyocyte differentiation, the resulting cell pellet was resuspended in ITS medium supplemented with CHIR 99021 (Axon Medchem, #1386) and BMP4 (R&D, 314-BP-10) based on differentiation efficiency screening (WT: 1.5 uM CHIR, 1.5 ng/mL BMP4 ; QKI KO: 1.25 uM CHIR, 1.5 ng/mL BMP4) (**Fig. 1a**), and seeded onto Matrigel-coated plates. To promote uniform cell distribution and adherence, the plates were gently agitated in a crosswise motion, tapped several times, and allowed to rest at room temperature for 20 minutes prior to incubation. After 24 hours, media was replaced with TS media (consisting of KO DMEM (Thermo Fisher Scientific, 10829018), 1% transferrin (Sigma Aldrich, T8158-1G), selenium (Sigma Aldrich Cat# S5261-10G), and 250 µM 2-Phospho-L-Ascorbic Acid (Sigma, 49752) and cultured for 24 hours, followed by replacement with TS media supplemented with 100nM C59 (Biomol, Cat# Cay16644) for 48 hours (D4 (cardiac progenitors) were crosslinked at this stage). Fresh media (TS+100 mM C59) was added, followed by 24 hours, followed by media changes with TS media every 48 hours. D8 (cardiomyocyte) samples were taken after confirmation of beating at days 7-8. For eCLIP, cells were UV crosslinked as previously described^49^. Briefly, cells were rinsed with PBS, a minimal amount of PBS was added to cover (∼2 mL per 10cm plate), followed by crosslinking (254 nM, 400 mJ/cm2) on ice. After crosslinking, cells were scraped and moved to conical tubes and pelleted, followed by removal of PBS and snap freezing. For RNA-seq, RNA was extracted using standard Trizol (Invitrogen) extraction.

### Cardioid differentiation

Cardiac organoids (cardioids) were generated using media and conditions adapted from a previous publication^97^. Briefly, hPSCs were grown in FTDA to approximately 70% confluency. For cardioid formation, 7500 cells per well were seeded into ultralow-attachment 96-well plates (Corning) and centrifuged for 5 min at 200g. After 24 hours, cells were induced with FLyABCH medium [Cardiac Differentiation Medium (CDM) containing FGF2 (30 ng/ml; Proteintech), activin A (50 ng/ml; Miltenyi Biotech), BMP4 (10 ng/ml; R&D Systems), 3 μM CHIR (Tocris), 5 μM LY294002 (Tocris), and insulin (1 μg/ml; Roche)] for 36 to 40 hours. Subsequently, cells were treated with BFIIWPRa medium [CDM, containing FGF2 (8 ng/ml), BMP4 (10 ng/ml), 1 μM IWR-1 (Tocris), and 0.5 μM retinoic acid (Sigma-Aldrich)] for 96 hours with medium change every 24 hours. Next, the medium was changed to BFI [CDM, containing BMP4 (10 ng/ml), FGF2 (8 ng/ml), and insulin (10 μg/ml)] for 48 hours with medium change after 24 hours. After 48 hours in BFI, cells were maintained in CDM+I (10 μg/ml of insulin) until they were harvested or used for imaging. For RNA isolation, three organoids were pooled per replicate and transferred into 250 μl of TRIzol solution. For imaging, organoids were fixed for 15 min in 4% paraformaldehyde (PFA), washed three times in PBS, and stored at 4°C until further processing.

### Generation of Stable Doxycycline-inducible QKI Overexpression Lines in QKI-KO hESCs

For the generation of stable, doxycycline-inducible rescue hESC lines, PiggyBac-based transposon-mediated genomic insertion was performed as previously described^95^. A total of 175,000 cells were seeded on Matrigel-coated 6-well dishes in FTDA growth medium supplemented with 10 µM Y-27632. The next day, a transfection mixture was prepared, consisting of 200 µl Opti-MEM, 12 µl Fugene, the desired insert (KA0717), transactivator (KA0637), and pBase plasmid in a 10:3:1 ratio, totaling 2.8 µg of DNA. After 15 minutes of incubation at room temperature, the medium was changed to fresh FTDA, and the transfection mixture was added dropwise.

Following a 24-hour incubation at 37°C, the medium was replaced with fresh FTDA supplemented with 0.5 µg/ml G-418 (Thermo Fisher Scientific, 11811023). Selection was maintained for 14 days in FTDA growth medium with G-418, including at least two passages. To verify proper integration and expression of the construct, cells were induced by adding 20 ng/ml doxycycline to the growth medium and incubated for 24 hours. The following day, GFP-positive cells were observed using a fluorescence microscope. For the cardiac differentiation of the resulting clones, 25-100 ng/ml doxycycline was added to the cardiac differentiation medium at various stages of the process.

### 293T and C2C12 Cell Culture and Transfections

HEK293T (Cat# 632180, Takara Bio) cells were cultured in high glucose Dulbecco’s Modified Eagle Medium (DMEM) (Fisher Scientific) supplemented with 10% Heat Inactivated Fetal Bovine Serum (FBS) (Thermo Fischer) and 100 ug/ml Penicillin Streptomycin (Thermo Fisher). C2C12 cells (gift from Dr. Thomas Cooper, BCM) were cultured in high glucose DMEM supplemented with 20% FBS and 100 ug/ml Pen/Strep. All cells were cultured in a humidified incubator at 37°C with 5% CO2.

For QKI siRNA experiments, HEK293T cells were seeded at a density of 3x10^5^ cells/ml in 6-well culture plates and incubated overnight, followed by transfection with either non-targeting siRNA (Silencer® select Negative control #1, Invitrogen) or siRNA against QKI (Silencer® select - ID s18084, Invitrogen) at a concentration of 30 µM per well using Lipofectamine RNAimax (Invitrogen) according to manufacturer’s guidelines. Cells were then incubated for 72 hr prior to collection.

For minigene assays, transfections of minigenes to assess target exon dependency on QKI were typically done in 12-well culture plates with an initial seeding density of 1x10^5^ cells/ml. The following morning, cells were transfected with either non-targeting siRNA (Silencer® select Negative control #1, Invitrogen) or siRNA against QKI (Silencer® select-ID s18084, Invitrogen) at a concentration of 1 µM per well using Lipofectamine™ RNAimax (Invitrogen) according to the manufacturer’s guidelines. On the evening of same day, the cells were subsequently transfected with 1 µg of minigene plasmid DNA (Viafect™, Promega) at a ratio of 1:2 (DNA: reagent) according to manufacturer’s guidelines, then incubated for 72 hr. The same transfection procedure was followed for C2C12 cells, with the exception that siRNA against mouse Qki (Silencer® select-ID n408900, Invitrogen) was used to knockdown endogenous Qki.

Myc-tagged QKI WT and KH mutant (K120A; R124A) expression plasmids were previously described^66^. For eCLIP studies, HEK293T cells were transiently transfected with 10 µg of either the WT or mutant plasmid in 10 cm plates (Viafect, Promega) at a ratio of 1:2 (DNA: reagent) according to manufacturer’s guidelines and incubated for 72 hr prior to UV crosslinking. Cells were UV crosslinked as previously described^49^. Briefly, cells were rinsed with PBS, a minimal amount of PBS was added to cover (∼2 mL per 10cm plate), followed by crosslinking (254 nM, 400 mJ/cm2) on ice. After crosslinking, cells were scraped and moved to conical tubes and pelleted, followed by removal of PBS and snap freezing.

### Profiling RNA interactions with eCLIP

QKI eCLIP from D0, D4, and D8 cardiomyocytes was performed using the original eCLIP method as previously described^49^, using RNA adapters X1A & X1B. Briefly, cells were lysed in iCLIP lysis buffer, followed by limited digestion with RNase I (Ambion), immunoprecipitation of RBP-RNA complexes with anti-QKI antibody (A300-183A, Fortis Life Sciences) and Protein G sheep anti-rabbit Dynabeads (ThermoFisher), and stringent washes. After dephosphorylation with FastAP (Thermo Fisher) and T4 PNK (NEB), a barcoded RNA adapter was ligated to the 3’ end (T4 RNA Ligase, NEB). Samples were then run on SDS-PAGE gels and transferred to nitrocellulose membranes, and a region from 40-115 kDa (75 kDa = 225 nt of RNA above the protein size) was isolated and proteinase K (NEB) treated to isolate RNA. RNA was reverse transcribed and treated with ExoSAP-IT (Affymetrix) to remove excess oligonucleotides. A second DNA adapter (containing a random-mer of 10 (N10) random bases at the 5’ end) was then ligated to the cDNA fragment 3’ end (T4 RNA Ligase, NEB). After cleanup (Dynabeads MyOne Silane, ThermoFisher), an aliquot of each sample was first subjected to qPCR (to identify the proper number of PCR cycles), and then the remainder was PCR amplified (Q5, NEB) and size selected via agarose gel electrophoresis.

HEK293T eCLIP experiments (including transgenic myc-tagged wild-type and mutant QKI) were performed using the seCLIP method as previously described^98^. As above, cells were lysed in iCLIP lysis buffer, followed by limited digestion with RNase I (Ambion) and immunoprecipitation of RBP-RNA complexes with either anti-MYC antibody (sc-40, Santa Cruz) or anti-QKI antibody (A300-183A, Fortis Life Sciences) as indicated along with Protein G sheep anti-rabbit Dynabeads (ThermoFisher), and stringent washes. Following library preparation was performed as previously described^98^, including SDS-PAGE gel electrophoresis, nitrocellulose membrane transfer and RNA isolation, reverse transcription (SuperScript IV, ThermoFisher), DNA adapter ligation containing a random-mer of 10 (N10) random bases at the 5’ end, and PCR amplification (Q5, NEB).

### eCLIP data analysis

Analyses of eCLIP sequencing data were performed using a previously described eCLIP processing pipeline^98^, with scripts available at https://github.com/VanNostrandLab/eclip. Briefly, 10nt of sequence was removed from the 5’ end of either read 2 (original eCLIP) or read 1 (seCLIP), followed by adapter trimming (cutadapt v1.14^99^), removal of reads mapping to repeat elements (STAR v2.5.1b^100^), and mapping to the human genome (hg19) (STAR v2.5.1b), with only uniquely mapping reads kept. PCR duplicates were then removed using either custom scripts (for original eCLIP) or umi_tools v1.1.1 (for seCLIP). Clusters of enriched reads were identified with CLIPPER^101^, with read density then compared against paired size-matched input datasets to identify input-enriched peaks as previously described^98^. Reproducible peaks were identified using a previously described framework^98^ based off of Irreproducible Discovery Rate (IDR)^51^ and modified here to incorporate three replicates for D0, D4, and D8 QKI eCLIP experiments. The association of peaks to genomic regions was performed as previously described^50^, based on the following priority order: tRNA, miRNA, miRNA-proximal (within 500 nt), CDS, 3′UTR, 5′UTR, 5′ splice site (within 100 nt of exon), 3′ splice site (within 100 nt of exon), proximal intron (within 500 nt of splice site region), distal intron (further than 500 nt from the splice site region), followed by noncoding exonic. Analysis of QKI and other RNA binding protein eCLIP data from K562 and HepG2 cells utilized previously described data from the ENCODE consortium^50^. For single-nucleotide resolution motif analysis, only the mapped position at the 5’ end of read 1 (seCLIP) or the 5’ end of read2 (original eCLIP) was utilized.

Mapping to U6 and other repetitive elements was performed based off of a previously described software pipeline^52^ available at https://github.com/VanNostrandLab/repetitive-element-mapping. Briefly, adapter-trimmed reads from above were mapped with bowtie2 (v2.2.6) against a custom database containing rRNA, small RNA (snoRNA, snRNA, vtRNA, yRNA, RN7SK, RN7SL, tRNA), simple repeats, and repetitive elements (RepBase v. 18.05)^102^ as previously described^52^. For this work, the underlying database of elements was modified to include N_10_ flanking each element, in order to better capture reads that overhang the end of included elements. As previously described, all equal scoring mappings were kept and compared against genomic mapping from the standard eCLIP pipeline above, with genomic mapping kept if it was more than 2 mismatches per read better than to the repeat element, and the repeat mapping kept otherwise. After PCR duplicate removal, quantification was then performed at both the individual element- and also RNA family-level to obtain final enrichments.

### RNA-seq library preparation

For cardiomyocytes, RNA was polyA-enriched, followed by library preparation with Illumina stranded mRNA kits. For 293T experiments, total RNA was extracted from cells using Trizol (Invitrogen) according to manufacturer guidelines followed by DNase (Zymo Research) treatment and clean-up using a Zymo RNA clean & Concentrator™-5 kit (Zymo Research). RNA concentration was determined using a Nanodrop and 3 µg of purified total RNA was used for Poly-A selection using the NEBNext® Poly(A) mRNA magnetic isolation module according to manufacturer’s guidelines. RNA-seq library preparation was performed utilizing the same protocol as seCLIP^98^, based off of a previously described RNA-seq approach^103^.

### RNA-seq gene expression and splicing analysis

For Hues6 RNA-seq data, adapter trimming was first performed (cutadapt v3.4), followed by removal of reads mapping (STAR v2.4.0j) to a database of repetitive elements, and finally mapping to human genome (hg19) with STAR (v2.4.0j), with only uniquely mapping reads kept. Gene expression was quantified by featureCounts (v2.0.1), followed by differential expression determined by DESeq2 (v1.30.0) analysis on protein-coding genes only. Alternative splicing was quantified with rmats-turbo (v4.3.0)^53^, with significant differential splicing events defined as those with rMATS change in Percent Spliced In |ΔPSI| ≥ 0.1, FDR ≤ 0.01, and at least 10 reads (exclusion + inclusion), with overlapping duplicates removed with the ‘subset_jxc’ tool as previously described^59^. 293T RNA-seq data was processed identically, except for initial removal of a 10nt UMI (removed and kept as a UMI from the 5’ end of read 1 and trimmed from the 3’ end of read 2) and its utilization to remove PCR duplicates after genome mapping (umi_tools v1.1.1). In both cases, to generate control sets of constitutive and ‘native SE’ cassette exons, junction-spanning reads (requiring at least 10nt overhanging both sides) were counted for all exon triplets in GENCODE v19, with PSI values determined for those with at least 30 junction-spanning reads. Constitutive exons were defined as those with no skipping observed in any wild-type replicate, whereas ‘native SE’ exons were defined as those with 0.5 ≤ PSI ≤ 0.95 in more than half of wild-type replicates. Integration of differential alternative splicing events with eCLIP data into splicing maps was performed with the rbp-maps software tool^59^ with default settings to normalize eCLIP IP minus input read density.

For further analyses, differential alternative cassette exons in D8 QKI KO versus wild-type RNA-seq were split into four classes based on their expression and splicing pattern in D0: “consistent” events with differential splicing in D0 (defined as D0 ΔPSI ≥ 0.1 for D8 QKI KO-excluded, or ΔPSI ≤ 0.1 for D8 QKI KO-included), “discordant” events expressed in D0 but which had no or opposite-direction QKI-dependent splicing (defined as D0 ΔPSI < 0 for D8 QKI KO-excluded, or ΔPSI > 0 for D8 QKI KO-included), “expression-driven” events that had low expression (less than 10 junction-spanning reads for exclusion and inclusion combined) in D0 wild-type, or “other”.

### Scoring of splice site strength

Maximum Enropy (MaxEnt) scores were determined using previously described software^61^. For U5- and U6-specific scores, the canonical 5’ splice site sequence GAGGUAAGUAU was utilized; the U5 score was defined as the number of positions exactly matching the -3 to -1 position “GAG”, whereas the U6 score was defined as the number of positions exactly matching the +4 to +8 position “AGUAU” sequence.

### Minigene constructs of QKI target exons

To construct minigene reporters, a region encompassing the QKI target exon and including immediate flanking introns and exons was amplified from HEK293T genomic DNA using Platinum™ SuperFi II Taq polymerase (Thermo Fisher Scientific) according to manufacturer’s guidelines in PCR fragments of 2 or 3 and assembled onto the backbone vector pcDNA 3.1 (+) (Thermo Fisher Scientific) using the NEBuilder HiFi DNA Assembly Master Mix kit (E2621L). In some cases, e.g for the RYR2 gene, parts of intronic regions were shortened (due to large intron lengths) to better facilitate cloning into the backbone vector. Assembled fragments were transformed into NEB^⍰^5-alpha competent cells (New England Biolabs, C2987H) according to manufacturer’s guidelines. 50-100 µl of transformed cells was spread onto LB selection plates containing 100 µg Ampicillin (Apex BioResearch) and incubated overnight at 37°C. Plasmid DNA was extracted from single clones using a Zymopure miniprep kit (Zymo Research) for mini-preparations and Purelink™ HiPure Plasmid Midiprep kit (Invitrogen) for midi-preparations. Sequence verification was done using whole plasmid sequencing by Plasmidsaurus utilizing Oxford Nanopore Technology with custom analysis and annotation. Construction of minigene mutants harboring 5’splice site mutations or QKI deletion motifs was done using site-directed mutagenesis and the original minigene construct as template for PCR amplification. Amplified fragments were assembled as previously described using the NEBuilder HiFi DNA Assembly Master Mix kit.

### RT-PCRs and Western Blots

After 72 hr of incubation, cells were collected by trypsinization and split into two halves in 1.5 ml microfuge tubes for either RNA or protein extraction. All cells were pelleted by centrifuging for 5 minutes at 1000 x g and supernatant was discarded. For RNA extraction, the pellet was resuspended in 300 ml TRIzol™ reagent (nvitrogen) and extracted according to manufacturer’s guidelines. RNA concentration was measured using a Nanodrop, after which 1 µg of total RNA was treated with Amplification grade DNAse (Invitrogen) according to manufacturer guidelines prior to cDNA synthesis. DNAse-free RNA was then used as template for reverse transcription using SuperScript™ IV First-Strand synthesis kit, according to manufacturer guidelines. Subsequently, 1 µl of cDNA was used in PCR reactions with Platinum™ SuperFi II Taq polymerase according to manufacturer guidelines and primer-specific annealing temperatures. PCR products were electrophoresed in 1X TBE (Tris-Borate EDTA) buffer and 1-1.5% Agarose gels stained with SybrSafe™ (Invitrogen) and visualized in an Azure imaging system (AZI300-01, Azure Biosystems).

Pellets for immunoblotting were resuspended in lysis buffer (50 mM Tri-HCl pH 7.4, 100 mM NaCl, 1% NP-40, 0.1% SDS 0.5% NaDC (Sodium deoxycholate) and incubated on ice for 5 minutes then centrifuged at 10, 000 x g for 10 minutes at 4°C. Cleared lysate (supernatant) was transferred to a clean microfuge tube and protein concentration determined using a Pierce protein BCA assay kit (Thermo Scientific). Approximately 10 µg of protein was mixed with 1X NuPAGE LDS sample buffer (Life Technologies) and run on precast 4-12% NuPAGE Bi-Tris gels (Invitrogen) with 1X MOPS SDS running buffer (Life Technologies). Protein was then transferred to PVDF membrane (Invitrogen) using an iBlot 2 transfer system (Invitrogen), blocked with Azure Chemiblot blocking buffer (Azure Biosystems) and incubated overnight with primary antibody at 4°C in a shaker. The next day the membrane was washed 3X for 5 minutes with 1X TBST (Boston Bioproducts) and incubated with secondary antibody. Immunoblots were probed for QKI knockdown using QKI antibody (Bethyl Laboratories) and either GAPDH (#51745, Cell Signaling Technology), Actin (#A5441, Millipore-SIGMA) or Tubulin (#3873S, Cell Signaling Technology) for loading controls. Unless otherwise stated, Trueblot anti-mouse IgG HRP (#18-8817-33, Rockland Immunochemicals), Trueblot anti-Rabbit IgG HRP (#18-8816-33, Rockland Immunochemicals) and anti-Rabbit IgG HRP (#7074S, Cell Signaling Technology) were used as secondary antibodies.

## NMR

### Protein expression and purification

The plasmid encoding the QKI^KH-Qua2^ (residue 76-205) sequence was cloned into a pET24 vector containing a His_6_-GB1 tag, and the recombinant protein was expressed in *E. coli* BL21 (DE3) cells in M9 minimal medium supplemented with 1 g/l ^15^NH_4_Cl. The bacterial cells were grown at 37 °C to an OD_600_ of 0.8, subsequently induced with 1.0 mM IPTG, and the protein was expressed at 20 °C overnight. The cells were harvested (7808 × *g*) and resuspended (50 mM Tris, pH 8.0, 500 mM NaCl, 10 mM imidazole, supplemented with lysozyme, 1 mg/ml DNase, 2 mM MgSO_4_, and protease inhibitor). After lysis of the cells using sonication and centrifugation (38759 × *g*, 1 h), the cleared lysate was added to Ni–NTA resin, washed with 2 M NaCl, and eluted with 500 mM imidazole. The His_6_-GB1 tag was cleaved with His-tagged TEV protease (4°C, overnight), and the protein was purified on a second Ni–NTA column. QKI^KH-Qua2^ was subsequently purified by ion-exchange chromatography on a HiTrap SP column (Cytiva) (20 mM Phos, pH 6.5, gradient from 0 to 1 M NaCl in 10 column volumes) followed by size-exclusion chromatography on a HiLoad 16/600 Superdex 75 column (GE Healthcare) (20 mM sodium phosphate, pH 6.5, 150 mM NaCl).

### In vitro transcription

13mer RNA, hairpin RNA, 32mer RNA was *in vitro* transcribed from a single-stranded DNA template containing the T7 promoter sequence (IDT Europe GmbH) using in-house produced 3 μl of T7 RNA Polymerase at 37 °C for 3 hours (0.64 μM DNA template, 0.64 μM T7 primer, 60 mM MgCl_2_, 8 mM of each rNTP, 5% PEG8000, 10% DMSO and 1 x transcription buffer). The reaction mix was purified on a DNAPac PA200 anion exchange column (22 x 250 mm, Thermo Fisher Scientific) by High-performance liquid chromatography (HPLC) (12.5 mM Tris, pH 8.0, 6 M Urea, gradient from 0 to 0.5 M sodium perchlorate) at 85 °C. Before use, the RNA was heated to 95 °C for 2 minutes and snap-cooled on ice.

### NMR spectroscopy

All NMR samples were measured at concentrations of 0.1 mM in NMR buffer (20 mM sodium phosphate, pH 6.5, 150 mM NaCl, 2 mM DTT) containing 10% (*v*/*v*) D_2_O at 25°C on 900-MHz Bruker Avance NMR spectrometers equipped with cryogenic triple-resonance gradient probes. All spectra were measured in 3 mm NMR tube. NMR spectra were processed with TOPSPIN3.5 (Bruker) and analyzed using NMRFAM-Sparky.^104^ Chemical shift assignments for QKI were transferred from published data.^105^

#### Titrations

^1^H,^15^N HSQC spectra were measured after each addition of titrant, and the changes were visualized by calculating the chemical shift perturbation (CSP) based on the following equation: 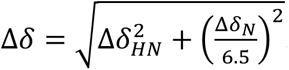.^106^ In the case of line-broadening of NMR signals the intensity of the peaks was used to assess the effect of binding.

### In Silico Modeling

The FASTA sequence of human QKI (Q96PU8) was fetched from the UniProt database^107^ for further structure modeling. Functional domains such as the homodimerization domain, the RNA-binding domain, and the KH domain were found using the family and domain category of the UniProt database. In particular, the domains found were Qua1 (residues 11-82), which promotes homodimerization; KH (residues 87-153); and Qua2 (residues 182-213), which participates in RNA binding. A BLAST^108^ search against the sequence found yielded a high hit with the RCSB PDB entry 4JVH^66^, an X-ray crystallography structure. This structure showed 60% query coverage and 96% identity to the query sequence, making it a good template for initial modeling.

To improve structural correctness, a sophisticated multi-template modeling with modeller software^109^ was used, based on three templates: 4JVH^66^, 4JVY^66^, and 2BL5^110^. These templates were chosen according to sequence similarity, structural alignment, and functional significance to the target protein. Combining multiple templates resolved structural ambiguities and enhanced model precision. Additional refinement was done with the help of prediction tools Robetta^111^ and I-TASSER^112^ that employ different algorithms like ab initio and hybrid modeling to yield high-confidence structural models. Of these models generated, the best-defined and most precise structure from the robust multi-template modeling method was chosen for all further analysis.

After generating the protein model through advanced modeling, the next step was to carry out protein-RNA docking to examine their interactions. U6 RNA was docked against monomeric and dimerized structures to see which conformation gives more stability to the complex. Since the generated QKI model happened to be a monomer, HADDOCK^113^ was utilized to generate a dimerized one.

For the docking experiments, residues around the U6 5’ stem loop and the 5’ splice site (5’ss) in the Pre-B (PDB ID: 6AH0^114^) and B (PDB ID: 5O9Z^115^) spliceosome structures were chosen. In addition to applying the U6 RNA structures from the Pre-B and B complexes, we also modeled the RNA with TrRosetta and RNA Composer. Of these, the RNA structure that we obtained with RNA Composer proved to be fuller. We therefore have three RNA structures docked against three different combinations of the protein domains for a full analysis of their interaction and stability.

Molecular Dynamics Simulation (MDS) allowed all the docked protein-RNA systems to be simulated. MDS was carried out on the three RNA structures, 2 from PDB IDs 6AH0, 5O9Z, and a modelled structure from RNA composer, with three combinations of protein domains in complex with each other using GROMACS (GROningen MAchine for Chemical simulations) software package, versions 2024.1^116^ and 2022.3^117^, but with the same forcefield.

The molecular dynamics simulations for all the complexes were performed using the AMBER99SB-ILDN^118^ force fields. Gromacs software was used to perform these molecular dynamics simulations. The individual complexes were solvated on the TIP3P model in a cubic or dodecahedron cell of 1.0 nm size. The topology files for the complexes were created using the Gromacs pdb2gmx command. A suitable balancing counter ion (Na^+^ or Cl^−^) was added to balance the system. Molecular dynamics simulations were conducted in three phases: energy minimization, equilibration, and production run. Steepest descent minimization was performed for a maximum of 50,000 steps with a tolerance of 1000 kJ/mol/nm. The Verlet cutoff scheme was employed, with a van der Waals cutoff of 1.2 nm and long-range electrostatics treated by the Particle mesh Ewald(PME) method^119^. NVT equilibration was performed for 50,000 steps with a time step of 1 fs using a Nose-Hoover^120^ thermostat at 300 K. This was succeeded by NPT equilibration employing a Berendsen barostat at 1 bar. LINCS algorithm^121^ held constraints in both instances. Parrinello-Rahman barostat^122^ was used to control pressure at 1 bar, while temperature was maintained at 300 K employing the Nose-Hoover thermostat. Molecular dynamics simulation was run until a stable root-mean-square deviation (RMSD) curve was attained to ensure equilibrium and reliable data for subsequent analysis. The final 50 ns of the stabilized trajectory were extracted and subjected to Molecular Mechanics Poisson-Boltzmann Surface Area (MMPBSA)^123^ analysis to evaluate the binding free energies of the complexes.

### Complexome Profiling

Sample preparation and high-resolution clear native electrophoresis (hrCNE) of hESCs were performed as previously described^124, 125^ with the following modifications. Cells were crosslinked using UVP CL-1000 Ultraviolet Crosslinker (254 nm, 400 mJ/cm²). Afterwards, cells were collected and lysed in Complete PB^124^ with 0.3% (v/v) NP-40 and 1x Halt Protease and Phosphatase Inhibitor Cocktail, EDTA-free. Nuclei were washed in Complete PB with 0.3% (v/v) NP-40 and resuspended in nuclear lysis buffer (20 mM HEPES, pH 7.4; 150 mM NaCl; 2.5 mM MgCl₂; 0.5 mM CaCl₂; 1% NP-40; 0.1 mM DTT; 1x Halt Protease and Phosphatase Inhibitor Cocktail, EDTA-free). The RNase-treated samples were mixed with RNase A (DNase and protease-free, 10 mg/ml, Thermo Scientific, Cat# EN0531), and incubated for 30 minutes at 22°C. The reaction was stopped by adding SUPERaseIn inhibitor. For samples without RNase treatment, SUPERaseIn was added directly during the lysis process.

Protein amount of 100 µg of each sample was supplemented with 5 µl 5% Coomassie G250 in 500 mM aminocaproic acid and 15 µl 0.1% Ponceau S in 50% glycerol. Samples were loaded on top of a 3-18% acrylamide gradient gel (dimensions 14x14 cm). After native electrophoresis in a cold chamber, blue-native gels were fixed in 50% (v/v) methanol, 10% (v/v) acetic acid, 10 mM ammonium acetate for 30 min and stained with Coomassie (0.025% Serva Blue G, 10% (v/v) acetic acid). Each lane of a BNE gel was cut into 48 equal fractions and collected in 96 filter well plates (30-40 µm PP/PE, Pall Corporation). The gel pieces were destained in 60% Methanol, 50 mM ammonium bicarbonate (ABC). Solutions were removed by centrifugation for 2 min at 600×g. Proteins were reduced in 10 mM DTT, 50 mM ABC for one hour at 56 °C, and alkylated for 45 min in 30 mM iodoacetamide. Samples were digested for 16 hours with trypsin (sequencing grade, Promega) at 37°C in 50 mM ABC, 0.01% Protease Max (Promega), and 1 mM CaCl2. Peptides were eluted in 30% acetonitrile and 3% formic acid, centrifuged into a fresh 96-well plate, dried in a speed vac, and resolved in 1% acetonitrile and 0.5% formic acid.

Liquid chromatography–mass spectrometry (LC-MS) analysis was performed using a Thermo Scientific Q Exactive HF mass spectrometer equipped with an easy nLC 1200 (Thermo Fisher Scientific, Bremen, Germany) fitted with a 35 cm, 75 μm ID fused-silica column packed in-house with 1.9-μm ReproSil-Pur C18 particles (Dr. Maisch). The column was maintained at 50 °C using an integrated column oven (Sonation). Peptides were eluted in a non-linear gradient of 5–28% ACN over 135 min and directly sprayed into the mass spectrometer using a nanoFlex ion source (Thermo Fisher Scientific). MS parameters were set to full-scan (400–1200 m/z) in profile mode at a resolution of 70,000 at m/z 200, a maximum injection time of 100 ms, and an AGC target value of 3 × 10E6. Up to 20 of the most intense peptides per full scan were isolated using a 1.5 Th window for fragmentation by higher-energy collisional dissociation (HCD) with a normalized collision energy (NCE) of 28. MS/MS spectra were acquired in centroid mode with a resolution of 17,500, a maximum injection time of 75 ms, and an AGC target value of 5 × 10^4^. Single charged ions, ions with a charge state > 6, and ions with unassigned charge states were not considered for fragmentation, and dynamic exclusion was set to 20 s to minimize the acquisition of fragment spectra representing already acquired precursors.

MS Data were analyzed by MaxQuant (v 2.6.7.0)^126^ using default settings. Proteins were identified using human reference proteome database UniProtKB with 82861 entries, released in 9/2024. The enzyme specificity was set to Trypsin. Acetylation (+42.01) at the N-terminus and oxidation of methionine (+15.99) were selected as variable modifications and carbamidomethylation (+57.02) as a fixed modification on cysteines. False discovery rate (FDR) for the identification of proteins and peptides was 1%. Intensity-based absolute quantification (IBAQ) values were recorded. The sum of all IBAQ values of data sets with RNAse A treatment was normalized to the proteins of the corresponding control without RNAse treatment. Protein abundance within native lanes was normalized to maximum appearance to enable comparison of complexes between samples.

### IP-Mass Spectrometry

Wild-type and QKI KO hESCs were crosslinked using UV light at 254 nm wavelength (UVP CL-1000 Ultraviolet Crosslinker, total output 400 mJ/cm²). Subsequently, cells were collected in lysis buffer (150 mM NaCl; 50 mM Tris, pH 7.5; 1% IGPAL-CA-630; 5% Glycerol; 1x Halt Protease and Phosphatase Inhibitor Cocktail, EDTA-free). RNase A (DNase- and protease-free, 10 mg/ml, Thermo Scientific, Cat# EN0531) was added, and the samples were incubated on ice for 20 minutes. RNase activity was halted by adding SUPERaseIn inhibitor. The lysate was clarified by centrifugation, and protein concentration was determined using a Detergent Compatible Bradford Assay Kit (Pierce, Cat# 23246). Dynabeads Protein G magnetic beads (Invitrogen. Cat# 10003) were equilibrated in lysis buffer (100 µl of slurry per 2 mg of total protein) and conjugated with anti-QKI antibody (A300-183A, Bethyl Laboratories; 10 µg per sample) for 30 minutes at 4°C on a rotator. The clarified lysate was incubated with the conjugated beads for 18 hours at 4°C on a rotator. Beads were then washed twice with wash buffer (150 mM NaCl; 50 mM Tris, pH 7.5) containing 0.05% IGPAL and five additional times with wash buffer.

Beads were resuspended in a lysis buffer containing 2% sodium deoxycholate (SDC), 1 mM tris(2-carboxyethyl)phosphine (TCEP), 4 mM chloroacetamide (CAA), and 50 mM Tris-HCl (pH 8.5), followed by incubation at 95 °C for 10 min. Protein digestion was performed overnight at 37 °C using Lys-C (Wako Chemicals) at a 1:50 enzyme-to-protein ratio (w/w) and trypsin (Promega, V5113) at a 1:100 ratio (w/w). Digestion was quenched by adding isopropanol/1% trifluoroacetic acid (TFA) to a final concentration of 75% (v/v). Peptides were purified using styrene-divinylbenzene reversed-phase sulfonate (SDB-RPS) material (Empore, 2241) and subsequently dried under vacuum. Dried peptides were reconstituted in 2% acetonitrile (ACN) / 0.1% formic acid (FA), and equal volumes were loaded onto an EASY-nLC 1200 system (Thermo Fisher Scientific, Bremen, Germany) coupled to a Q Exactive HF hybrid quadrupole-Orbitrap mass spectrometer (Thermo Fisher Scientific, Bremen, Germany).

Peptide separation was performed on a 35 cm × 75 μm inner diameter fused-silica column packed with 1.9 μm C18 particles (ReproSil-Pur, Dr. Maisch), maintained at 50 °C using an integrated column oven (Sonation). Chromatographic separation was achieved using a linear gradient from 4% to 38% solvent B over 60 min, followed by an increase to 60% B in 5 min, and 95% B in 1 min, which was maintained for an additional 5 min. Solvent A consisted of 0.1% FA in water, and solvent B of 0.1% FA in 80% ACN.

MS data were acquired in data-dependent acquisition (DDA) mode. Full MS scans were recorded at a resolution of 60,000 (at *m/z* 200) over an *m/z* range of 300–1,650, with a maximum injection time of 20 ms and an automatic gain control (AGC) target of 3 × 10⁶. The top 10 most intense precursors were selected for higher-energy collisional dissociation (HCD) with a normalized collision energy (NCE) of 27 and isolated using a 1.4 Th quadrupole window.

MS/MS spectra were acquired in centroid mode at a resolution of 30,000 (at *m/z* 200), with a maximum injection time of 54 ms and an AGC target of 1 × 10⁵. Dynamic exclusion was set to 20 s to minimize repeated sequencing of identical precursors.

Raw data files were processed using MSFragger v4.1 via FragPipe v22.0, incorporating IonQuant v1.10.27 and Philosopher v4.3. The default FragPipe workflow was used, with “Normalize intensity across runs” and “Add MaxLFQ” disabled. Database searches were performed against the *Homo sapiens* reference proteome from UniProt (ID: UP000005640, downloaded February 2025). Trypsin was specified as the protease, allowing up to two missed cleavages. The precursor mass tolerance and fragment mass tolerance were both set to ±20 ppm. Cysteine carbamidomethylation was defined as a fixed modification, while methionine oxidation and protein N-terminal acetylation were set as variable modifications. Protein quantification was based on label-free quantification (LFQ) intensities from IonQuant. Identifications flagged as contaminants or lacking quantification were removed. Intensities were median-normalized within replicates, and missing values were imputed using global minimum replacement. Statistical analysis was performed using an empirical Bayes moderated t-test implemented in the limma R package (Ritchie et al., 2015). Benjamini-Hochberg false discovery rate (FDR) correction was applied, and proteins with an adjusted p-value ≤ 0.05 and an absolute log₂ fold-change ≥ 1 were considered significantly regulated.

### Co-immunoprecipitation of QKI and Spliceosome proteins

To assess endogenous QKI interaction with tri-snRNP proteins, co-immunoprecipitation assays were performed using HEK293 cell lysates. Briefly for each sample, cells were seeded at a density of 2x10^6^ in 10 cm or 5x10^6^ in 15 cm culture plates, at least 72 hr prior to co-immunoprecipitation assays. Confluent plates were washed twice in 1X PBS on ice and lysed in non-denaturing co-IP lysis buffer (20 mM Tris-HCl pH 8, 137 mM NaCl, 10% Glycerol, 1% Nonidet P-40 (NP-40), 2 mM EDTA) supplemented with 1X protease inhibitor cocktail III (Cat# 539134, Millipore/Sigma). Cells were then scraped onto a 1.5 ml pre-cooled microfuge and incubated on ice for 15 minutes. The lysate was subsequently sonicated using a QSonica Bioruptor (Q800R3, QSonica) at 20% amplitude for 10 sec on a 2 sec ON and 5 sec OFF setting followed by centrifugation at 14,000 x g in a pre-cooled centrifuge for 10 minutes. During cell lysis incubation, beads were coupled to antibodies as follows; 125 µl per sample of sheep anti-rabbit Dynabeads M-280 (Cat# 11204D, Life Technologies) was washed twice in co-IP lysis buffer and incubated with 10 µg/sample (10 cm plate) or 15 µg/sample (15 cm plate) of either QKI (Cat# A300-183A), RBM42 (Cat# A305-138A, Bethyl Laboratories), SNRNP27 (Cat# HPA034541, Sigma-Aldrich) or TXNL4A (Cat# PA5-120069, Thermo Fischer Scientific) antibodies for 45 minutes at room temperature on a rotator. Following incubation, antibody-bead complexes were washed twice in lysis buffer and resuspended in either 100 µl (10 cm) or 125 µl (15 cm) lysis buffer. Supernatants from cleared lysates at the previous step were transferred to a clean microfuge tube and 2% of the lysate was aliquoted and stored as input. Antibody-beads complexes were then added to cleared lysates, and the mixture incubated overnight at 4°C in a rotator. Samples for IgG isotype controls (Rabbit IgG; Cat# 02-1602, Thermo Fischer Scientific) were processed in the same manner as antibody samples and incubated in parallel. The next day samples were briefly centrifuged at 2500 x g for 30 seconds and the supernatant removed. The beads were then washed up to six times with co-IP wash buffer (0.5% Triton-X-100) and resuspended in 2X SDS loading buffer, boiled at 90°C for 5 minutes and run on a 4-12% SDS PAGE followed by immunoblotting.

---

**Extended Data Fig. 1:**
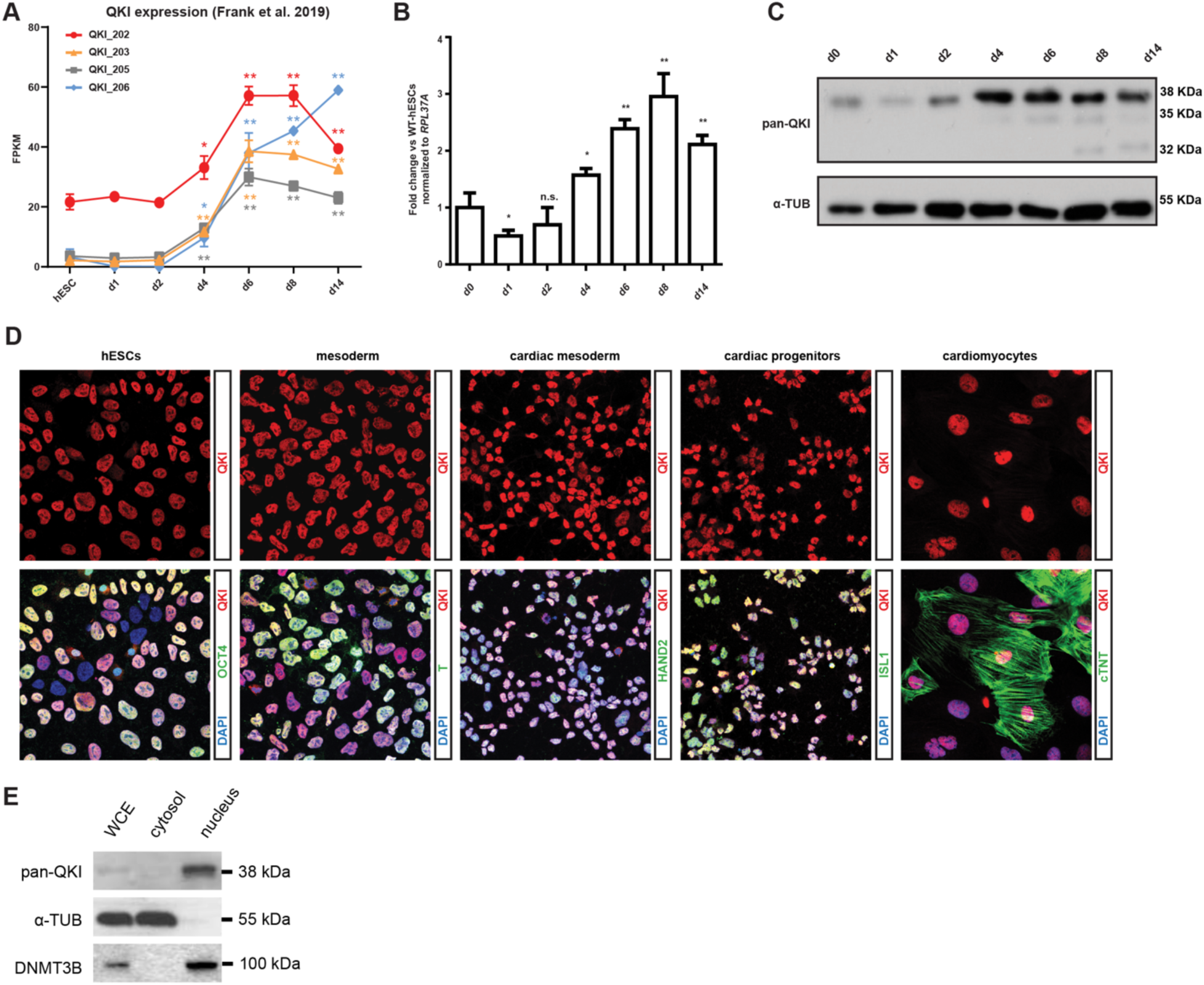
QKI is dynamically regulated and predominantly nuclear localized during cardiac differentiation of hESCs. **a**, RNA-seq quantification of individual QKI isoforms (QKI-202, -203, -205, -206) across cardiac differentiation stages (FPKM values from Frank et al., 2019), showing dynamic isoform expression patterns. **b**, qRT–PCR analysis of total QKI transcript levels at seven key time points during directed cardiac differentiation of hESCs, showing stage-resolved dynamic regulation. Data represent mean ± SEM from *n* = 3 biological replicates. **c**, Immunoblot of total QKI protein during cardiac differentiation from D0 to D14, confirming stage-specific regulation. α-Tubulin serves as a loading control. **d**, Immunofluorescence staining for QKI and stage-specific lineage markers at matched differentiation stages: OCT4 (pluripotent hESCs), T (primitive streak/mesoderm), HAND2 (cardiac mesoderm), ISL1 (cardiac progenitors), and cTnT (cardiomyocytes). QKI expression is observed throughout all stages and localizes near-exclusively to the nucleus. **e**, Western blot analysis of nuclear and cytosolic fractions in D8 cardiomyocytes, showing enrichment of QKI in the nuclear compartment. DNMT3B and α-Tubulin serve as nuclear and cytosolic markers, respectively. Error bars represent ±SEM; *p-*values are calculated using Student’s *t*-test; biological replicates *n* = 3

**Extended Data Fig. 2:**
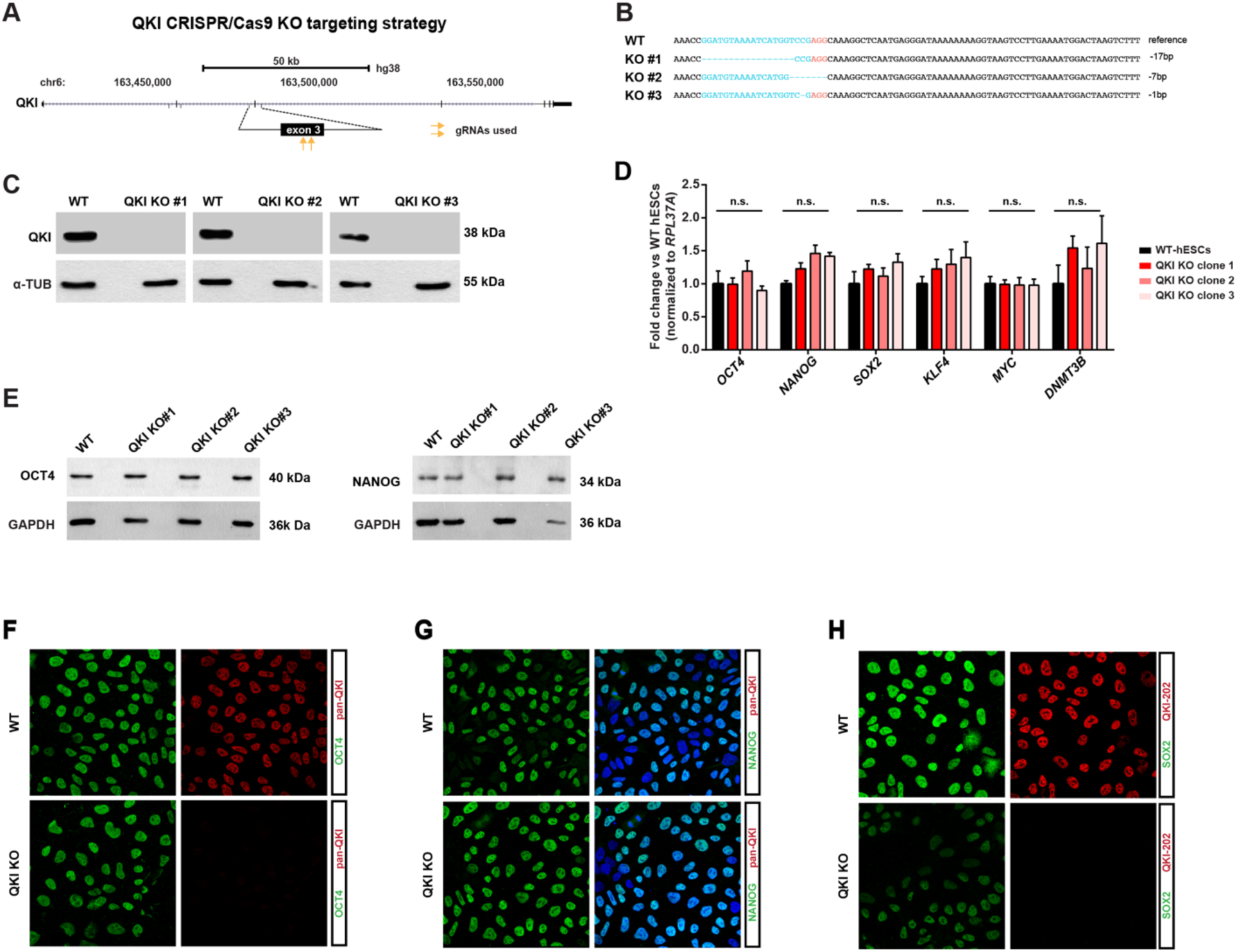
Generation and validation of QKI knockout hESCs. **a**, Schematic of the genomic QKI locus in humans along with the CRISPR/Cas9 strategy targeting exon 3 of QKI. **b**, Illustration of QKI knockout alleles in the indicated hESC clones, each harboring unique homozygous frameshift mutations disrupting the open reading frame of QKI. **c**, Immunoblot confirming loss of QKI protein in three independent knockout clones; α-Tubulin serves as loading control. **d**, qPCR validation of pluripotency markers (*OCT4, NANOG, SOX2, KLF4, MYC, DNMT3B*) showing no significant differences between WT and QKI KO hESCs clones (normalized to *RPL37A*). **e,** Immunoblot confirming the expression of pluripotency markers OCT4 and NANOG in QKI KO clones, comparable to the isogenic WT counterpart. **f**,**g,h**, Immunofluorescence analysis of pluripotency markers in WT and QKI KO hESCs. Representative images showing OCT4, NANOG, and SOX2 expression in wild-type and QKI knockout hESCs. Note the reduced SOX2 protein levels in QKI knockout cells compared to wild type, despite intact expression of OCT4 and NANOG. Error bars represent ±SEM; *p-*values are calculated using Student’s *t*-test; biological replicates *n* = 3

**Extended Data Fig. 3:**
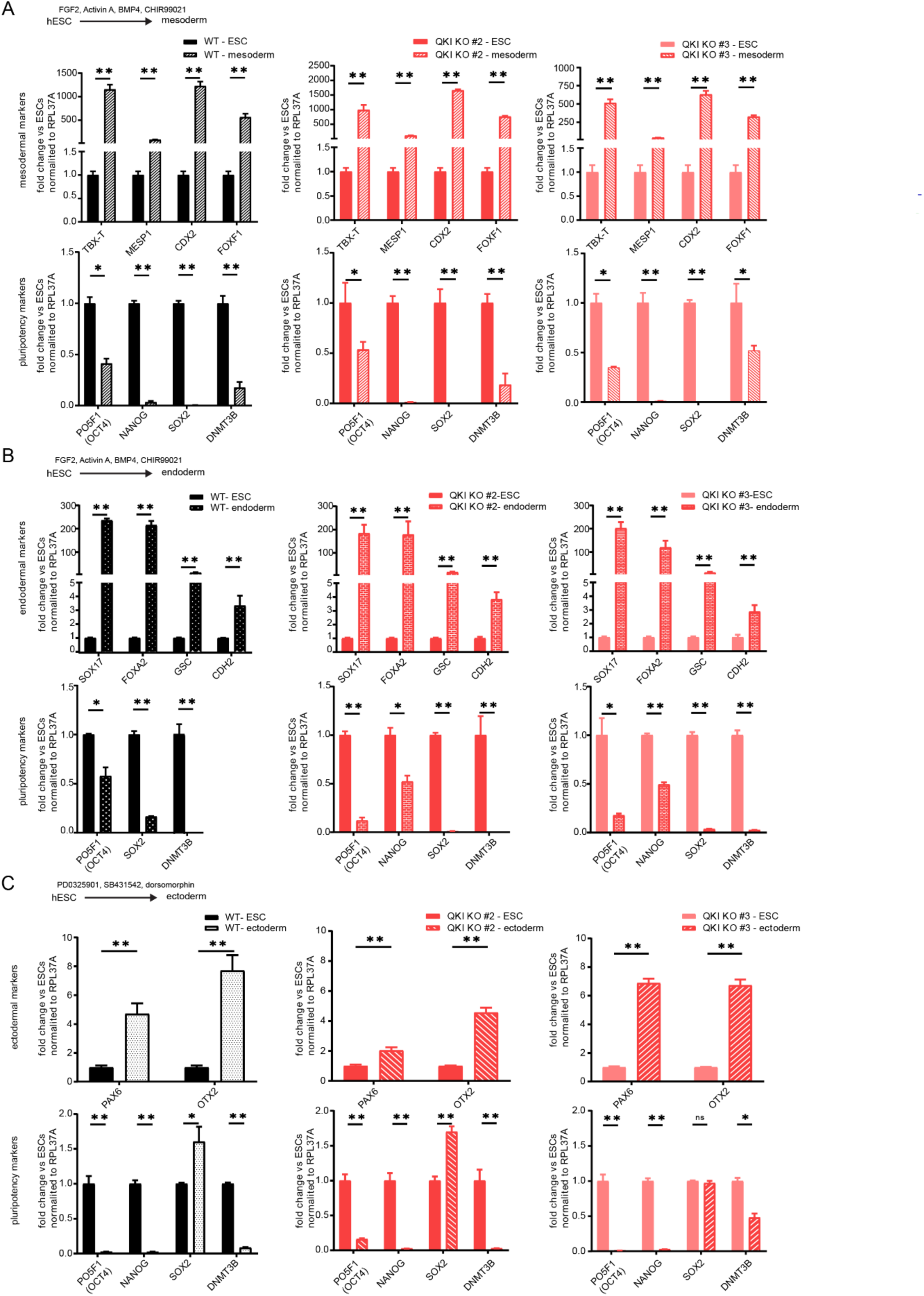
QKI knockout does not impair trilineage differentiation. **a**, Schematic of the directed mesoderm differentiation protocol. Below, qRT-PCR analysis of mesoderm markers (*TBXT, MESP1, CDX2, FOXF1*) in WT and three independent QKI KO hESC clones. All KO clones exhibit robust induction of mesoderm markers, indicating that QKI loss does not impair mesoderm commitment. **b**, Schematic of the directed endoderm induction protocol. qRT-PCR analysis of definitive endoderm markers (*SOX17, FOXA2, GSC, CDH2*) following directed endoderm differentiation of WT and QKI KO hESCs. All three KO clones show comparable induction of endodermal genes, demonstrating intact endodermal potential in the absence of QKI. **c**, Schematic of the directed ectoderm induction protocol. qRT-PCR quantification of ectoderm lineage markers (*PAX6, OTX2*) during directed ectoderm differentiation. All QKI KO clones activate canonical ectoderm genes to levels similar to WT, confirming that QKI is dispensable for ectoderm specification. Error bars represent ±SEM; *p-*values are calculated using Student’s *t*-test; biological replicates *n* = 3

**Extended Data Fig. 4:**
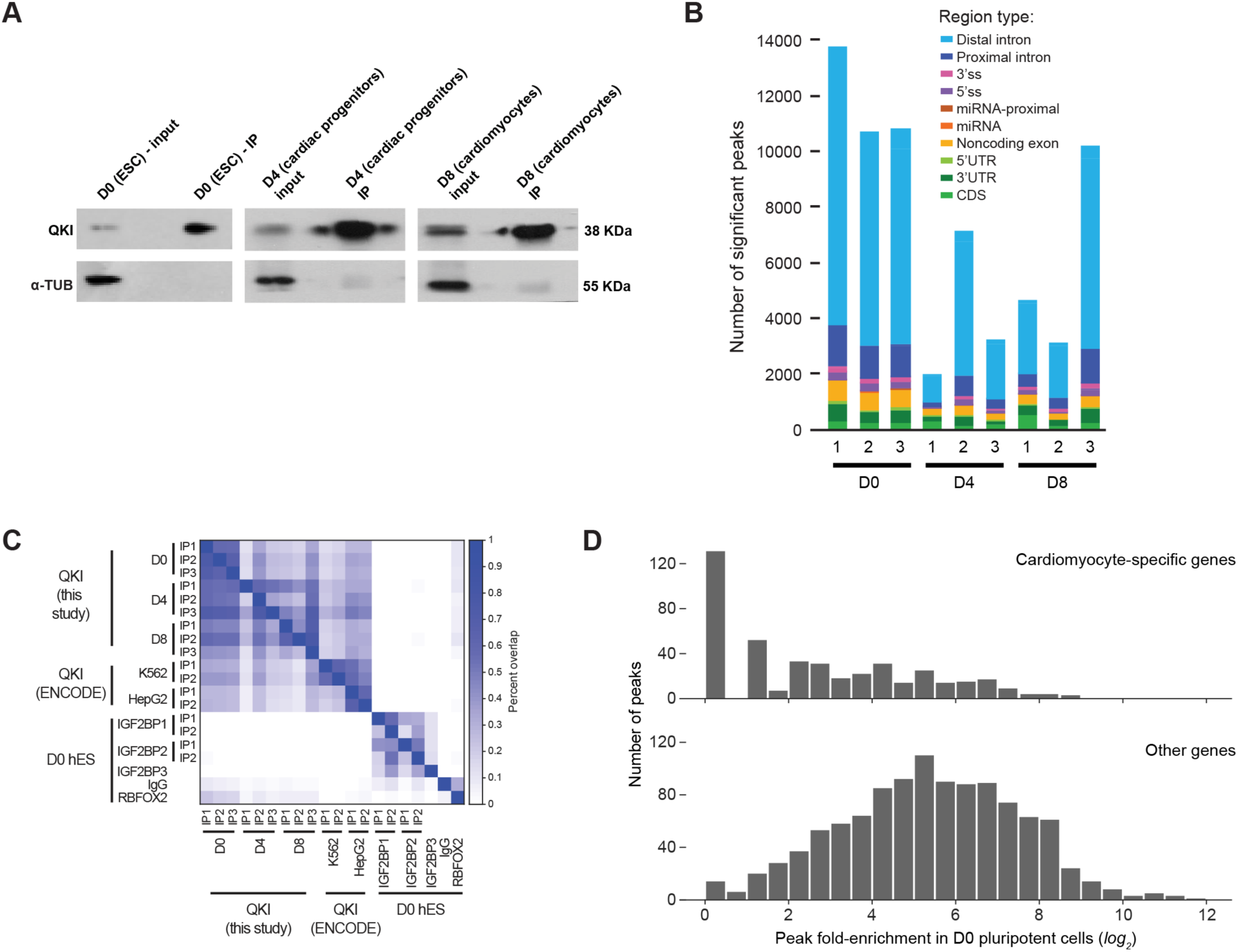
Validation and specificity of QKI eCLIP libraries. **a**, Immunoblot confirming successful immunoprecipitation (IP) of endogenous QKI from lysates under eCLIP conditions from hESCs (D0), cardiac progenitor (D4), and cardiomyocyte (D8) stages. α-Tubulin serves as loading control. **b**, Bars indicate fraction of significant peaks (*p* ≤ 10^-3^, fold-enrichment ≥ 8) from individual replicate eCLIP experiments, which overlap indicated RNA region annotations. **c**, Clustering analysis comparing QKI eCLIP replicates from this study across differentiation stages with publicly available QKI eCLIP datasets (ENCODE) and unrelated RBPs (IGF2BP1, IGF2BP3, and RBFOX2). QKI replicates cluster tightly together and separates clearly from unrelated RBPs, confirming both the quality and target specificity of QKI eCLIP across stages. Additionally, stage-resolved QKI datasets exhibit progressive differences, indicating developmental regulation of QKI binding. **d**, Histogram of peak fold-enrichment in D0 QKI eCLIP for peaks identified in D8 QKI eCLIP (as shown in Fig. 2i). Genes were separated as ‘cardiomyocyte-specific’ (five-fold or higher expression in D8 than D0) or ‘other genes’.

**Extended Data Fig. 5:**
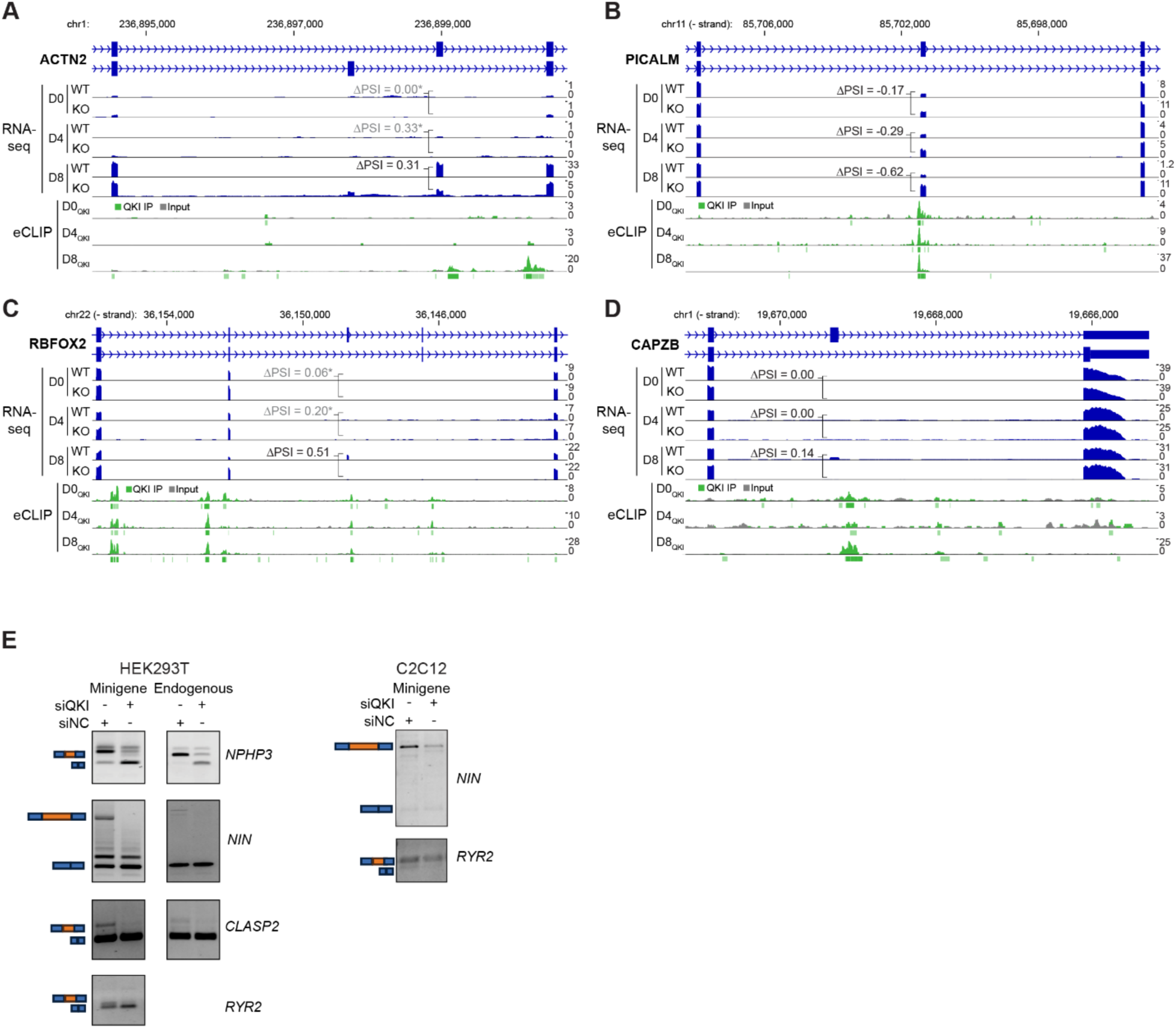
QKI loss leads to nuclear sequestration of mis-spliced cardiac transcripts. **a-d**, Representative read density tracks showing RNA-seq and QKI eCLIP signal for differentially spliced exons in *ACTN2, PICALM, RBFOX2,* and *CAPZB* across cardiac differentiation. **e**, Representative RT-PCR assays to validate splicing defects for indicated QKI-dependent targets using (left) minigene reporters and corresponding endogenous transcripts (*NPHP3, NIN, CLASP2,* and *RYR2*) in HEK293T cells following QKI knockdown, or (right) C2C12 cells.

**Extended Data Fig. 6:**
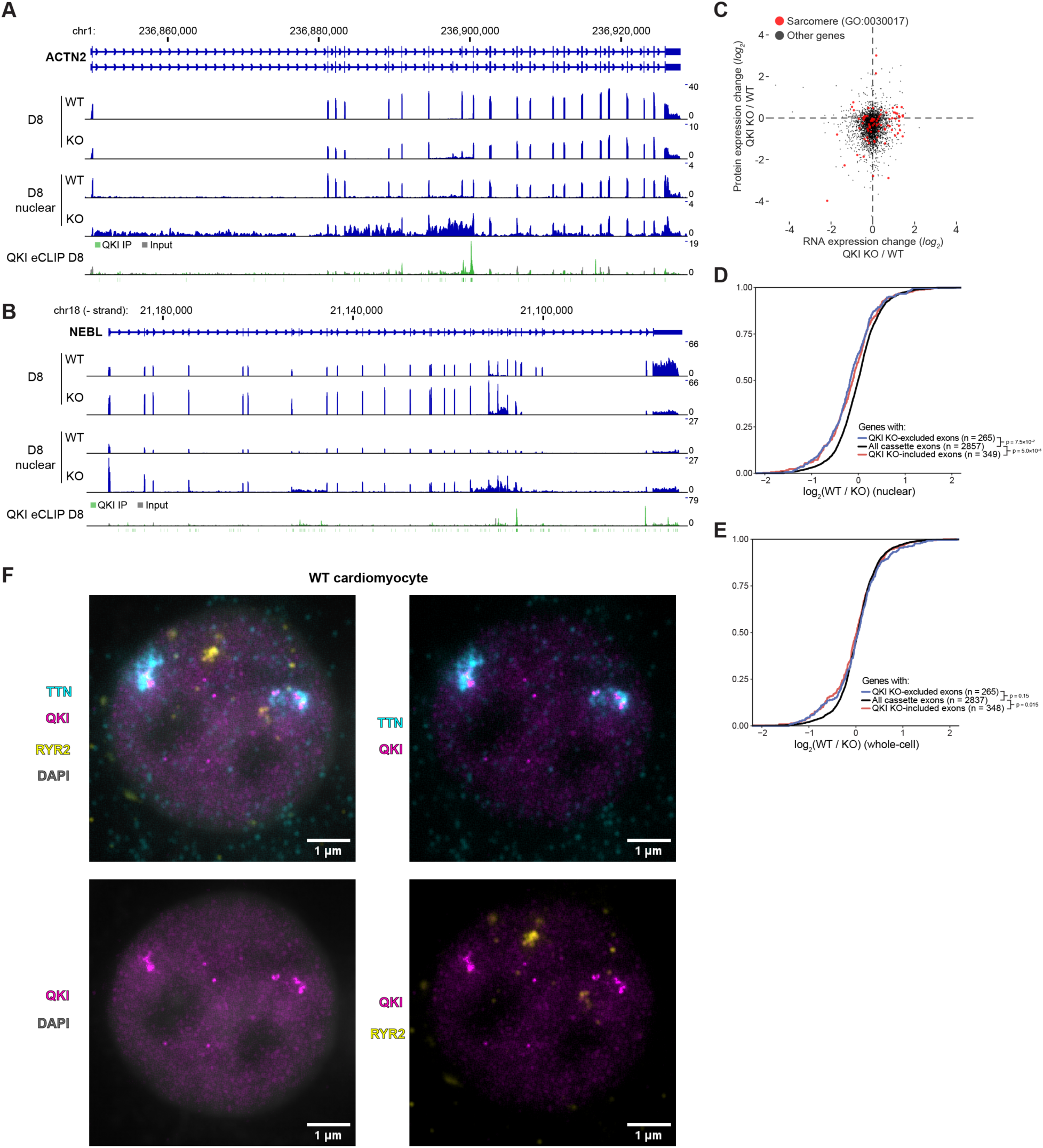
QKI knockdown leads to mis-splicing and nuclear retention of RNA for essential cardiac factors. **a-b**, Whole-gene read density plots for (**a**) *ACTN2* and (**b**) *NEBL*, including QKI eCLIP (and paired input), D0, D4, and D8 polyA RNA-seq, and D8 nuclear fraction rRNA-depleted RNA-seq. **c**, Scatter plot indicates lack of correlation between RNA and protein expression changes in D8 QKI knockout versus wild-type cardiomyocytes. Sarcomere genes (Gene Ontology ID: 0030017) are indicated in red. **d-e**, Distribution of indicated transcript categories in (**d**) nuclear and (**e**) whole-cell RNA-seq between WT and QKI KO cardiomyocytes. Transcripts containing QKI KO–excluded exons show modest but significant nuclear enrichment compared to all cassette exons (P = 7.5 × 10⁻⁷, two-sided Wilcoxon rank-sum test), consistent with the nuclear retention of mis-spliced RNAs. **f**, Maximum intensity projections of TTN and RYR2 smRNA FISH combined with immunofluorescence for QKI in human cardiomyocytes. *TTN* transcripts form discrete nuclear foci that wrap around QKI-enriched puncta, suggestive of localized accumulation at *TTN* transcription sites. In contrast*, RYR2* transcripts form distinct foci that do not colocalize with either *TTN* or QKI, indicating that QKI–*TTN* condensates do not serve as general hubs for all QKI-dependent transcripts. This spatial segregation underscores the transcript-specific nature of the nuclear interactions of QKI in cardiomyocytes.

**Extended Data Fig. 7:**
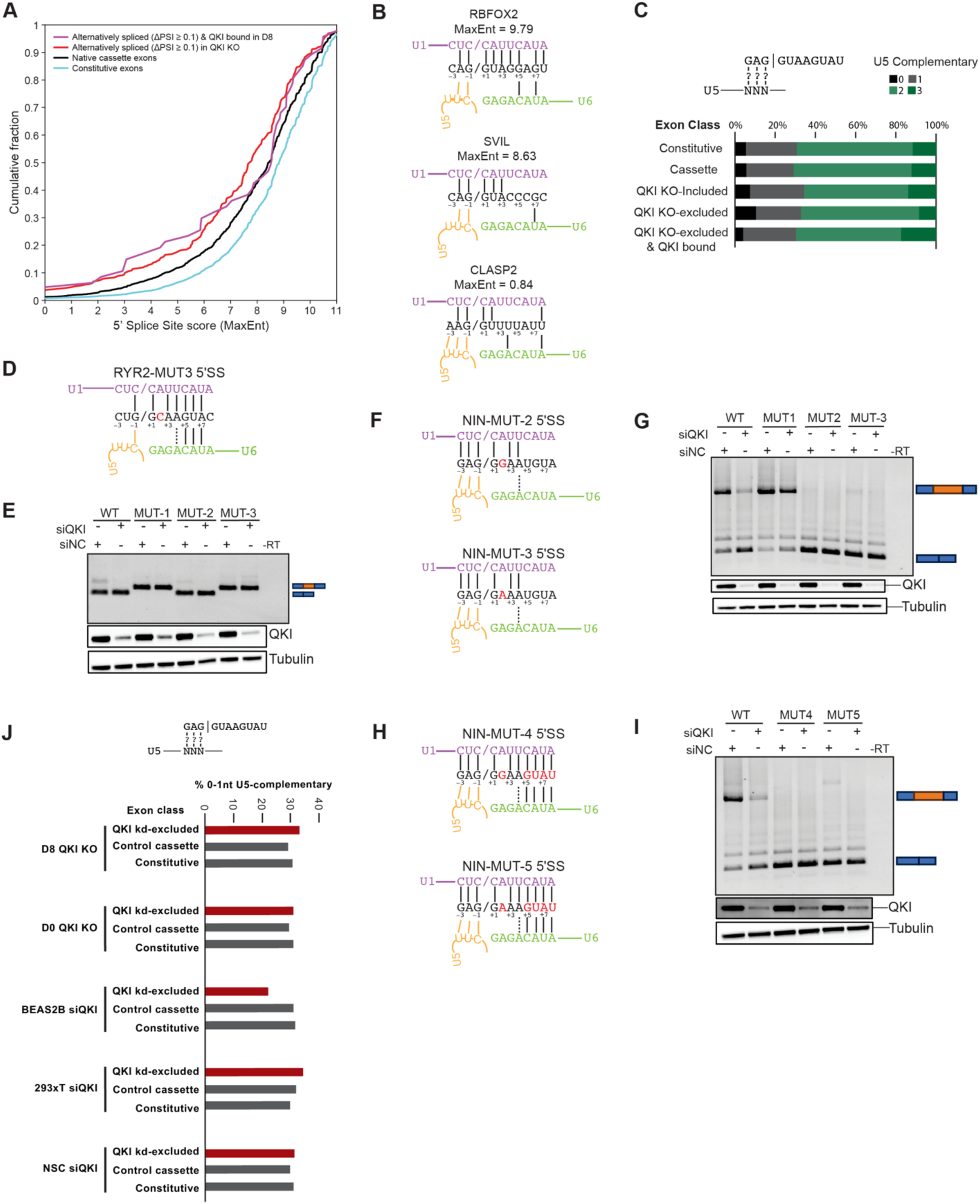
U6 complementarity determines QKI dependency at weak 5ʹSS. **a**, MaxEnt 5ʹSS strength scores of all cassette exons, QKI-regulated cassette exons (ΔPSI ≥ 0.1, FDR ≤ 0.01), and QKI-regulated exons bound by QKI with strong inclusion (ΔPSI ≥ 0.5), revealing a significant shift toward weaker 5ʹSS in QKI-dependent targets. **b**, Schematic of 5ʹSS with indicated base-pairing potential to U1, U5, and U6 snRNAs for exons in cardiac-related genes that lose exon inclusion upon QKI knockout. **c**, For indicated exon classes, bars show the fraction of exons with their 5ʹSS containing the indicated number of complementary nucleotides to U5 snRNA. **d**, Schematic of the RYR2 MUT3 minigene, which harbors a +2C mutation in the context of the inherent good complementarity to U6 snRNA in the *RYR2* 5ʹSS. **e**, Representative RT-PCR image of the *RYR2 MUT3* minigene splicing profile relative to WT, MUT1, and MUT2 minigenes that have been described previously. *RYR2* MUT3 results in consistent inclusion of exon 75, similar to MUT1, indicating a context-specific significance for specific nucleotides at +2 in QKI-dependent exon inclusion. **f**, Schematic of NIN minigenes harboring a *+*2T>G (MUT2) and a *+*2T>A (MUT3) 5ʹSS mutations to test the significance of nucleotide identity +2T in the *NIN* 5ʹSS context. **g**, RT-PCR image showing splicing profiles of *NIN* MUT2 and MUT3 relative to WT and MUT1 minigenes. In this context, altering the nucleotide identity at +2 leads to a complete loss of inclusion in a QKI-independent manner. **h**, As in **f** but with splice sites harboring mutations that improve complementarity to U6 snRNA. **i**, RT-PCR showing comparative splicing profiles of *NIN* MUT5 and MUT6 to the *NIN* WT minigene. Improving complementarity to U6 snRNA, in this context, does not enhance inclusion of *NIN* exon 18, indicating a context-specific requirement for +2T in the NIN 5ʹSS. **j**, As in Fig. 5j, predicted U5 complementarity was scored for QKI knockout- or knockdown-excluded exons from our profiling of D8 and D0 QKI knockout cells, our profiling of QKI siRNA knockdown in 293T cells, and published QKI knockdown in *BEAS2B* and neuronal stem cell (NSC) cells^60, 64^, with no enrichment seen for weak U5 complementarity.

**Extended Data Fig. 8:**
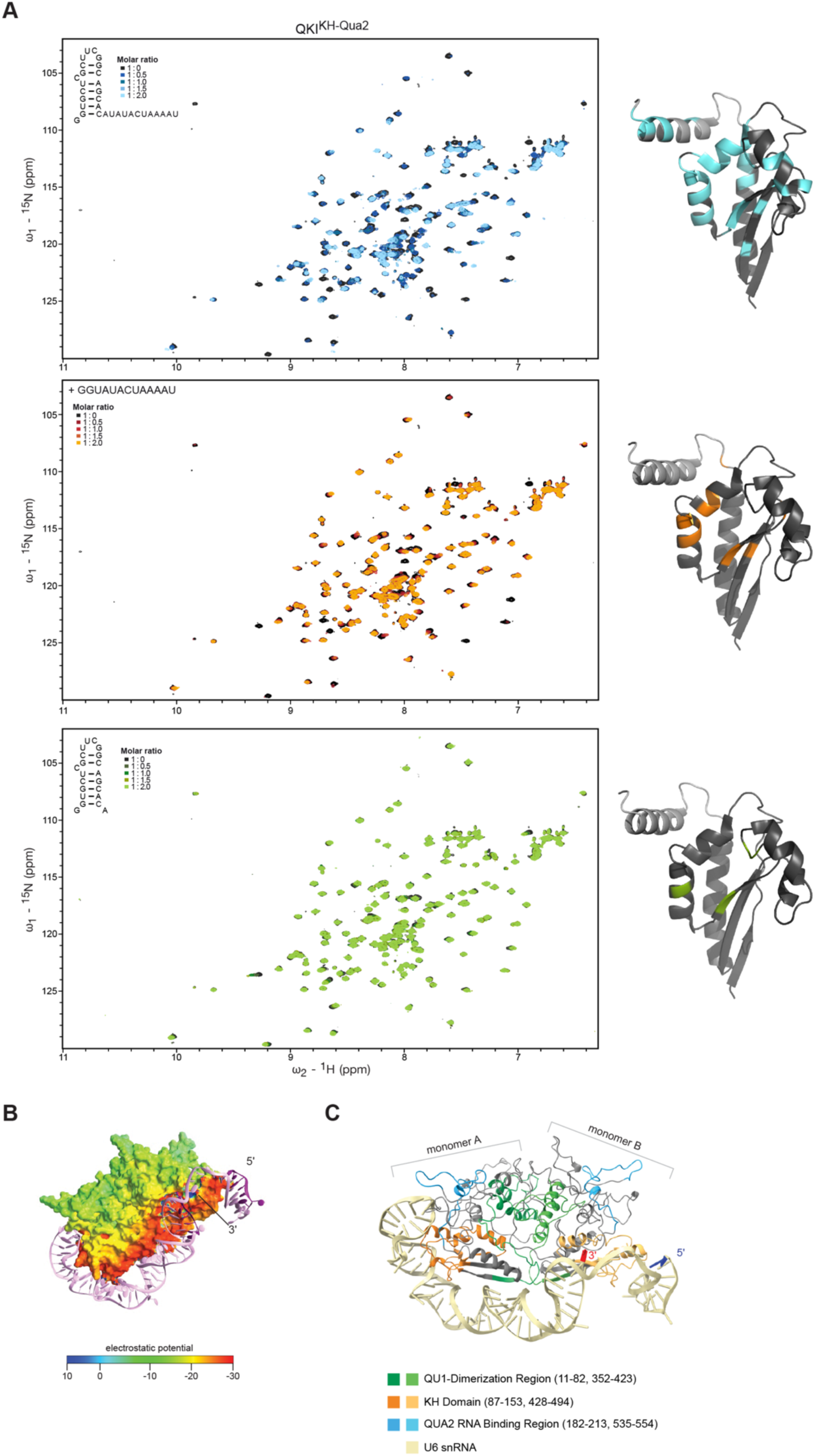
Molecular basis of QKI–U6 snRNA interaction. **a**, Representative 1H‒15N HSQC NMR titrations of recombinant 15N-labeled QKI KH‒QUA2 domain with increasing concentrations of U6 RNA. Chemical shift perturbations confirm specific interaction with the 5ʹ region of U6, consistent with structured RNA recognition. Amino acid residues interacting with the RNA molecules (disappearing peaks and peaks with chemical shift perturbation > 0.05) are highlighted in the structural models (pdb: 4jvh). **b,** Molecular docking model showing the 3D arrangement of U6 snRNA interacting with the QKI dimer. U6 wraps around QKI, positioning its 5ʹ end along the positively charged RNA-binding groove, consistent with the binding footprint revealed by NMR. **c,** Electrostatic surface potential of QKI dimer highlighting a positively charged groove that accommodates U6 snRNA. The U6 backbone follows a trajectory consistent with the modeled RNA-binding interface.

## REFERENCES

1. Marasco, L.E. & Kornblihtt, A.R. The physiology of alternative splicing. Nature reviews. Molecular cell biology 24, 242–254 (2023).

2. Rogalska, M.E., Vivori, C. & Valcarcel, J. Regulation of pre-mRNA splicing: roles in physiology and disease, and therapeutic prospects. Nature reviews. Genetics 24, 251–269 (2023).

3. Barbosa-Morais, N.L. et al. The evolutionary landscape of alternative splicing in vertebrate species. Science 338, 1587–1593 (2012).

4. Ule, J. & Blencowe, B.J. Alternative Splicing Regulatory Networks: Functions, Mechanisms, and Evolution. Molecular cell 76, 329–345 (2019).

5. Baralle, F.E. & Giudice, J. Alternative splicing as a regulator of development and tissue identity. Nature reviews. Molecular cell biology 18, 437–451 (2017).

6. Mazin, P.V., Khaitovich, P., Cardoso-Moreira, M. & Kaessmann, H. Alternative splicing during mammalian organ development. Nature genetics 53, 925–934 (2021).

7. Olthof, A.M., White, A.K. & Kanadia, R.N. The emerging significance of splicing in vertebrate development. Development 149 (2022).

8. Chen, J. et al. Differential alternative splicing landscape identifies potentially functional RNA binding proteins in early embryonic development in mammals. iScience 27, 109104 (2024).

9. Gotthardt, M. et al. Cardiac splicing as a diagnostic and therapeutic target. Nature reviews. Cardiology 20, 517–530 (2023).

10. Giudice, J. et al. Alternative splicing regulates vesicular trafficking genes in cardiomyocytes during postnatal heart development. Nature communications 5, 3603 (2014).

11. Gomes-Silva, B. et al. Alternative splicing dynamics during human cardiac development *in vivo* and *in vitro*. bioRxiv, 2025.2003.2021.642423 (2025).

12. Mazin, P. et al. Widespread splicing changes in human brain development and aging. Molecular systems biology 9, 633 (2013).

13. Recinos, Y. et al. Lineage-specific splicing regulation of MAPT gene in the primate brain. Cell Genom 4, 100563 (2024).

14. Li, Z. et al. RNA splicing controls organ-wide maturation of postnatal heart in mice. Developmental cell 60, 236–252 e238 (2025).

15. van den Hoogenhof, M.M., Pinto, Y.M. & Creemers, E.E. RNA Splicing: Regulation and Dysregulation in the Heart. Circulation research 118, 454–468 (2016).

16. Weyn-Vanhentenryck, S.M. et al. Precise temporal regulation of alternative splicing during neural development. Nature communications 9, 2189 (2018).

17. Li, C. et al. Alternative splicing categorizes organ development by stage and reveals unique human splicing variants linked to neuromuscular disorders. The Journal of biological chemistry 301, 108542 (2025).

18. Wang, E.T. et al. Alternative isoform regulation in human tissue transcriptomes. Nature 456, 470–476 (2008).

19. Black, D.L. Mechanisms of alternative pre-messenger RNA splicing. Annu Rev Biochem 72, 291–336 (2003).

20. Fu, X.D. & Ares, M., Jr. Context-dependent control of alternative splicing by RNA-binding proteins. Nature reviews. Genetics 15, 689–701 (2014).

21. Capitanchik, C., Wilkins, O.G., Wagner, N., Gagneur, J. & Ule, J. From computational models of the splicing code to regulatory mechanisms and therapeutic implications. Nature reviews. Genetics 26, 171–190 (2025).

22. Martinez-Lumbreras, S., Morguet, C. & Sattler, M. Dynamic interactions drive early spliceosome assembly. Curr Opin Struct Biol 88, 102907 (2024).

23. Wilkinson, M.E., Charenton, C. & Nagai, K. RNA Splicing by the Spliceosome. Annu Rev Biochem 89, 359–388 (2020).

24. Wahl, M.C., Will, C.L. & Luhrmann, R. The spliceosome: design principles of a dynamic RNP machine. Cell 136, 701–718 (2009).

25. Garcia-Ruiz, S. et al. Splicing accuracy varies across human introns, tissues, age and disease. Nature communications 16, 1068 (2025).

26. Taliaferro, J.M., et al. RNA Sequence Context Effects Measured In Vitro Predict In Vivo Protein Binding and Regulation. Molecular cell 64, 294–306 (2016).

27. Mao, M. et al. Modeling and Predicting the Activities of Trans-Acting Splicing Factors with Machine Learning. Cell systems 7, 510–520 e514 (2018).

28. Lee, Y. & Rio, D.C. Mechanisms and Regulation of Alternative Pre-mRNA Splicing. Annu Rev Biochem 84, 291–323 (2015).

29. Wong, M.S., Kinney, J.B. & Krainer, A.R. Quantitative Activity Profile and Context Dependence of All Human 5’ Splice Sites. Molecular cell 71, 1012–1026 e1013 (2018).

30. Roca, X., Krainer, A.R. & Eperon, I.C. Pick one, but be quick: 5’ splice sites and the problems of too many choices. Genes Dev 27, 129–144 (2013).

31. Yeo, G.W., Van Nostrand, E., Holste, D., Poggio, T. & Burge, C.B. Identification and analysis of alternative splicing events conserved in human and mouse. Proc Natl Acad Sci U S A 102, 2850–2855 (2005).

32. Parker, M.T., Fica, S.M. & Simpson, G.G. RNA splicing: a split consensus reveals two major 5’ splice site classes. Open Biol 15, 240293 (2025).

33. Montanes-Agudo, P., Pinto, Y.M. & Creemers, E.E. Splicing factors in the heart: Uncovering shared and unique targets. Journal of molecular and cellular cardiology 179, 72–79 (2023).

34. Chen, X. et al. QKI is a critical pre-mRNA alternative splicing regulator of cardiac myofibrillogenesis and contractile function. Nature communications 12, 89 (2021).

35. Fagg, W.S. et al. Definition of germ layer cell lineage alternative splicing programs reveals a critical role for Quaking in specifying cardiac cell fate. Nucleic acids research 50, 5313–5334 (2022).

36. Montanes-Agudo, P. et al. The RNA-binding protein QKI governs a muscle-specific alternative splicing program that shapes the contractile function of cardiomyocytes. Cardiovascular research 119, 1161–1174 (2023).

37. Bonnet, A. et al. Quaking RNA-Binding Proteins Control Early Myofibril Formation by Modulating Tropomyosin. Developmental cell 42, 527–541 e524 (2017).

38. de Bruin, R.G. et al. Quaking promotes monocyte differentiation into pro-atherogenic macrophages by controlling pre-mRNA splicing and gene expression. Nature communications 7, 10846 (2016).

39. Zhao, Z. et al. QKI shuttles internal m(7)G-modified transcripts into stress granules and modulates mRNA metabolism. Cell 186, 3208–3226 e3227 (2023).

40. Lee, J. et al. QUAKING Regulates Microexon Alternative Splicing of the Rho GTPase Pathway and Controls Microglia Homeostasis. Cell reports 33, 108560 (2020).

41. Sakers, K. et al. Loss of Quaking RNA binding protein disrupts the expression of genes associated with astrocyte maturation in mouse brain. Nature communications 12, 1537 (2021).

42. Hardy, R.J. et al. Neural cell type-specific expression of QKI proteins is altered in quakingviable mutant mice. The Journal of neuroscience : the official journal of the Society for Neuroscience 16, 7941–7949 (1996).

43. Frank, S. et al. yylncT Defines a Class of Divergently Transcribed lncRNAs and Safeguards the T-mediated Mesodermal Commitment of Human PSCs. Cell stem cell 24, 318–327 e318 (2019).

44. Bartsch, D. et al. Translational specialization in pluripotency by RBPMS poises future lineage-decisions. bioRxiv, 2021.2004.2012.439420 (2021).

45. Rao, J. et al. Stepwise Clearance of Repressive Roadblocks Drives Cardiac Induction in Human ESCs. Cell stem cell 18, 554–556 (2016).

46. Hofbauer, P. et al. Cardioids reveal self-organizing principles of human cardiogenesis. bioRxiv, 2020.2007.2006.189431 (2020).

47. Galarneau, A. & Richard, S. Target RNA motif and target mRNAs of the Quaking STAR protein. Nat Struct Mol Biol 12, 691–698 (2005).

48. Galarneau, A. & Richard, S. The STAR RNA binding proteins GLD-1, QKI, SAM68 and SLM-2 bind bipartite RNA motifs. BMC Mol Biol 10, 47 (2009).

49. Van Nostrand, E.L. et al. Robust transcriptome-wide discovery of RNA-binding protein binding sites with enhanced CLIP (eCLIP). Nat Methods 13, 508–514 (2016).

50. Van Nostrand, E.L. et al. A large-scale binding and functional map of human RNA-binding proteins. Nature 583, 711–719 (2020).

51. Li, Q., Brown, J.B., Huang, H. & J., B.P. Measuring reproducibility of high-throughput experiments. The Annals of Applied Statistics 5, 1752–1779 (2011).

52. Van Nostrand, E.L. et al. Principles of RNA processing from analysis of enhanced CLIP maps for 150 RNA binding proteins. Genome Biol 21, 90 (2020).

53. Wang, Y. et al. rMATS-turbo: an efficient and flexible computational tool for alternative splicing analysis of large-scale RNA-seq data. Nat Protoc 19, 1083–1104 (2024).

54. Zhang, X. et al. Cell-Type-Specific Alternative Splicing Governs Cell Fate in the Developing Cerebral Cortex. Cell 166, 1147–1162 e1115 (2016).

55. Hayakawa-Yano, Y. & Yano, M. An RNA Switch of a Large Exon of Ninein Is Regulated by the Neural Stem Cell Specific-RNA Binding Protein, Qki5. Int J Mol Sci 20 (2019).

56. Tharp, C.A., Haywood, M.E., Sbaizero, O., Taylor, M.R.G. & Mestroni, L. The Giant Protein Titin’s Role in Cardiomyopathy: Genetic, Transcriptional, and Post-translational Modifications of TTN and Their Contribution to Cardiac Disease. Front Physiol 10, 1436 (2019).

57. Kania, E.E., et al. Nascent transcript O-MAP reveals the molecular architecture of a single-locus subnuclear compartment built by RBM20 and the TTN RNA. bioRxiv (2024).

58. Witten, J.T. & Ule, J. Understanding splicing regulation through RNA splicing maps. Trends Genet 27, 89–97 (2011).

59. Yee, B.A., Pratt, G.A., Graveley, B.R., Van Nostrand, E.L. & Yeo, G.W. RBP-Maps enables robust generation of splicing regulatory maps. RNA 25, 193–204 (2019).

60. Hayakawa-Yano, Y. et al. An RNA-binding protein, Qki5, regulates embryonic neural stem cells through pre-mRNA processing in cell adhesion signaling. Genes Dev 31, 1910–1925 (2017).

61. Yeo, G. & Burge, C.B. Maximum entropy modeling of short sequence motifs with applications to RNA splicing signals. J Comput Biol 11, 377–394 (2004).

62. Parker, M.T. et al. m(6)A modification of U6 snRNA modulates usage of two major classes of pre-mRNA 5’ splice site. Elife 11 (2022).

63. Ishigami, Y., Ohira, T., Isokawa, Y., Suzuki, Y. & Suzuki, T. A single m(6)A modification in U6 snRNA diversifies exon sequence at the 5’ splice site. Nature communications 12, 3244 (2021).

64. Zong, F.Y. et al. The RNA-binding protein QKI suppresses cancer-associated aberrant splicing. PLoS Genet 10, e1004289 (2014).

65. Rappsilber, J., Ajuh, P., Lamond, A.I. & Mann, M. SPF30 is an essential human splicing factor required for assembly of the U4/U5/U6 tri-small nuclear ribonucleoprotein into the spliceosome. The Journal of biological chemistry 276, 31142–31150 (2001).

66. Teplova, M. et al. Structure-function studies of STAR family Quaking proteins bound to their in vivo RNA target sites. Genes Dev 27, 928–940 (2013).

67. Staley, J.P. & Guthrie, C. An RNA switch at the 5’ splice site requires ATP and the DEAD box protein Prp28p. Mol Cell 3, 55–64 (1999).

68. Charenton, C., Wilkinson, M.E. & Nagai, K. Mechanism of 5’ splice site transfer for human spliceosome activation. Science 364, 362–367 (2019).

69. Schmitzova, J., Cretu, C., Dienemann, C., Urlaub, H. & Pena, V. Structural basis of catalytic activation in human splicing. Nature 617, 842–850 (2023).

70. Agafonov, D.E. et al. Molecular architecture of the human U4/U6.U5 tri-snRNP. Science 351, 1416–1420 (2016).

71. Keiper, S. et al. Smu1 and RED are required for activation of spliceosomal B complexes assembled on short introns. Nature communications 10, 3639 (2019).

72. Prieto-Garcia, C. et al. Pathogenic proteotoxicity of cryptic splicing is alleviated by ubiquitination and ER-phagy. Science 386, 768–776 (2024).

73. Zhang, Z. et al. Cryo-EM analyses of dimerized spliceosomes provide new insights into the functions of B complex proteins. EMBO J 43, 1065–1088 (2024).

74. Zhang, Z. et al. Structural insights into the cross-exon to cross-intron spliceosome switch. Nature 630, 1012–1019 (2024).

75. Makarova, O.V., Makarov, E.M. & Luhrmann, R. The 65 and 110 kDa SR-related proteins of the U4/U6.U5 tri-snRNP are essential for the assembly of mature spliceosomes. The EMBO journal 20, 2553–2563 (2001).

76. Karaduman, R., Chanarat, S., Pfander, B. & Jentsch, S. Error-Prone Splicing Controlled by the Ubiquitin Relative Hub1. Molecular cell 67, 423–432 e424 (2017).

77. White, D.S., Dunyak, B.M., Vaillancourt, F.H. & Hoskins, A.A. A sequential binding mechanism for 5’ splice site recognition and modulation for the human U1 snRNP. Nature communications 15, 8776 (2024).

78. Rogalska, M.E. et al. Transcriptome-wide splicing network reveals specialized regulatory functions of the core spliceosome. Science 386, 551–560 (2024).

79. Sibley, C.R., Blazquez, L. & Ule, J. Lessons from non-canonical splicing. Nature reviews. Genetics 17, 407–421 (2016).

80. Artzt, K. & Wu, J.I. STAR trek: An introduction to STAR family proteins and review of quaking (QKI). Advances in experimental medicine and biology 693, 1–24 (2010).

81. Chen, X. et al. The Emerging Roles of the RNA Binding Protein QKI in Cardiovascular Development and Function. Front Cell Dev Biol 9, 668659 (2021).

82. Yu, Y. & Reed, R. FUS functions in coupling transcription to splicing by mediating an interaction between RNAP II and U1 snRNP. Proceedings of the National Academy of Sciences of the United States of America 112, 8608–8613 (2015).

83. Jutzi, D. et al. Aberrant interaction of FUS with the U1 snRNA provides a molecular mechanism of FUS induced amyotrophic lateral sclerosis. Nature communications 11, 6341 (2020).

84. Lopez, A.J. Alternative splicing of pre-mRNA: developmental consequences and mechanisms of regulation. Annu Rev Genet 32, 279–305 (1998).

85. Huang, S.C. et al. RBFOX2 promotes protein 4.1R exon 16 selection via U1 snRNP recruitment. Mol Cell Biol 32, 513–526 (2012).

86. Campagne, S. et al. Molecular basis of RNA-binding and autoregulation by the cancer-associated splicing factor RBM39. Nat Commun 14, 5366 (2023).

87. Kenny, C.J. et al. LUC7 proteins define two major classes of 5’ splice sites in animals and plants. Nature communications 16, 1574 (2025).

88. Zahler, A.M. et al. SNRP-27, the C. elegans homolog of the tri-snRNP 27K protein, has a role in 5’ splice site positioning in the spliceosome. RNA 24, 1314–1325 (2018).

89. Parker, M.T., Fica, S.M., Barton, G.J. & Simpson, G.G. Inter-species association mapping links splice site evolution to METTL16 and SNRNP27K. Elife 12 (2023).

90. Marian, A.J. & Braunwald, E. Hypertrophic Cardiomyopathy: Genetics, Pathogenesis, Clinical Manifestations, Diagnosis, and Therapy. Circulation research 121, 749–770 (2017).

91. Kong, S.W. et al. Heart failure-associated changes in RNA splicing of sarcomere genes. Circ Cardiovasc Genet 3, 138–146 (2010).

92. Kornienko, J. et al. Mislocalization of pathogenic RBM20 variants in dilated cardiomyopathy is caused by loss-of-interaction with Transportin-3. Nature communications 14, 4312 (2023).

93. Maatz, H. et al. RNA-binding protein RBM20 represses splicing to orchestrate cardiac pre-mRNA processing. The Journal of clinical investigation 124, 3419–3430 (2014).

94. Guo, W. et al. RBM20, a gene for hereditary cardiomyopathy, regulates titin splicing. Nature medicine 18, 766–773 (2012).

95. Rao, J. et al. Stepwise Clearance of Repressive Roadblocks Drives Cardiac Induction in Human ESCs. Cell Stem Cell 18, 341–353 (2016).

96. Zhang, M. et al. Universal cardiac induction of human pluripotent stem cells in two and three-dimensional formats: implications for in vitro maturation. Stem Cells 33, 1456–1469 (2015).

97. Hofbauer, P. et al. Cardioids reveal self-organizing principles of human cardiogenesis. Cell 184, 3299–3317 e3222 (2021).

98. Blue, S.M. et al. Transcriptome-wide identification of RNA-binding protein binding sites using seCLIP-seq. Nat Protoc 17, 1223–1265 (2022).

99. Martin, M. Cutadapt removes adapter sequences from high-throughput sequencing reads. EMBnet.journal 17, 10–12 (2011).

100. Dobin, A. et al. STAR: ultrafast universal RNA-seq aligner. Bioinformatics 29, 15–21 (2013).

101. Lovci, M.T. et al. Rbfox proteins regulate alternative mRNA splicing through evolutionarily conserved RNA bridges. Nat Struct Mol Biol 20, 1434–1442 (2013).

102. Bao, W., Kojima, K.K. & Kohany, O. Repbase Update, a database of repetitive elements in eukaryotic genomes. Mob DNA 6, 11 (2015).

103. Shishkin, A.A. et al. Simultaneous generation of many RNA-seq libraries in a single reaction. Nat Methods 12, 323–325 (2015).

104. Lee, W., Tonelli, M. & Markley, J.L. NMRFAM-SPARKY: enhanced software for biomolecular NMR spectroscopy. Bioinformatics 31, 1325–1327 (2015).

105. Maguire, M.L. et al. Solution Structure and Backbone Dynamics of the KH-QUA2 Region of the Xenopus STAR/GSG Quaking Protein. Journal of Molecular Biology 348, 265–279 (2005).

106. Mulder, F.A., Schipper, D., Bott, R. & Boelens, R. Altered flexibility in the substrate-binding site of related native and engineered high-alkaline Bacillus subtilisins. J Mol Biol 292, 111–123 (1999).

107. UniProt, C. UniProt: the Universal Protein Knowledgebase in 2025. Nucleic Acids Res 53, D609–D617 (2025).

108. Madden, T. The BLAST sequence analysis tool, in The NCBI Handbook, 2nd *edition.* 425–436 (2013).

109. Sali, A. & Blundell, T.L. Comparative protein modelling by satisfaction of spatial restraints. J Mol Biol 234, 779–815 (1993).

110. Maguire, M.L. et al. Solution structure and backbone dynamics of the KH-QUA2 region of the Xenopus STAR/GSG quaking protein. J Mol Biol 348, 265–279 (2005).

111. Kim, D.E., Chivian, D. & Baker, D. Protein structure prediction and analysis using the Robetta server. Nucleic Acids Res 32, W526–531 (2004).

112. Zhang, Y. I-TASSER server for protein 3D structure prediction. BMC Bioinformatics 9, 40 (2008).

113. de Vries, S.J., van Dijk, M. & Bonvin, A.M. The HADDOCK web server for data-driven biomolecular docking. Nat Protoc 5, 883–897 (2010).

114. Zhan, X., Yan, C., Zhang, X., Lei, J. & Shi, Y. Structures of the human pre-catalytic spliceosome and its precursor spliceosome. Cell Res 28, 1129–1140 (2018).

115. Bertram, K. et al. Cryo-EM Structure of a Pre-catalytic Human Spliceosome Primed for Activation. Cell 170, 701–713 e711 (2017).

116. Abraham, M. et al. (Zenodo; 2024).

117. Bauer, P., Hess, B. & Lindahl, E. (Zenodo; 2023).

118. Lindorff-Larsen, K. et al. Improved side-chain torsion potentials for the Amber ff99SB protein force field. Proteins 78, 1950–1958 (2010).

119. Essmann, U. et al. A smooth particle mesh Ewald method. The Journal of Chemical Physics 103, 8577–8593 (1995).

120. Evans, D.J. & Holian, B.L. The Nose–Hoover thermostat. The Journal of Chemical Physics 83, 4069–4074 (1985).

121. Hess, B., Bekker, H., Berendsen, H.J.C. & Fraaije, J.G.E.M. LINCS: A linear constraint solver for molecular simulations. Journal of Computational Chemistry 18, 1463–1472 (1997).

122. Parrinello, M. & Rahman, A. Polymorphic transitions in single crystals: A new molecular dynamics method. Journal of Applied Physics 52, 7182–7190 (1981).

123. Genheden, S. & Ryde, U. The MM/PBSA and MM/GBSA methods to estimate ligand-binding affinities. Expert Opin Drug Discov 10, 449–461 (2015).

124. Melnik, S. et al. Isolation of the protein and RNA content of active sites of transcription from mammalian cells. Nat Protoc 11, 553–565 (2016).

125. Wittig, I., Braun, H.P. & Schagger, H. Blue native PAGE. Nat Protoc 1, 418–428 (2006).

126. Cox, J. & Mann, M. MaxQuant enables high peptide identification rates, individualized p.p.b.-range mass accuracies and proteome-wide protein quantification. Nat Biotechnol 26, 1367–1372 (2008).

